# Spatial Pattern Analysis using Closest Events (SPACE) – A Nearest Neighbor Point Pattern Analysis Framework for Assessing Spatial Relationships from Image Data

**DOI:** 10.1101/2023.05.17.541131

**Authors:** Andrew M. Soltisz, Peter F. Craigmile, Rengasayee Veeraraghavan

**Author notes:** Co-corresponding authors Co-corresponding Author Contact Information Andrew M. Soltisz Mailing Address: 1028 Ponderosa Way, Woodland Park, Colorado 80863, United States ORCID: 0000-0002-1493-3544 Rengasayee Veeraraghavan Mailing Address: The Ohio State University, Institute for Behavioral Medicine Research, Room 415A, 460 Medical Center Dr, Columbus, Ohio 43210-1229, United States ORCID: 0000-0002-8364-2222. **Author Permanent Addresses** Andrew M. Soltisz 1028 Ponderosa Way, Woodland Park, Colorado 80863, United States Peter F. Craigmile The Ohio State University, College of Arts and Sciences, Department of Statistics, Cockins Hall, Room 404, 1958 Neil Ave, Columbus, Ohio 43210, United States Rengasayee Veeraraghavan The Ohio State University, College of Engineering, Department of Biomedical Engineering, Institute for Behavioral Medicine Research, Room 415A, 460 Medical Center Dr, Columbus, Ohio 43210-1229, United States. **<u>Declarations:</u>** Competing Interests The authors declare none. **Intellectual Property** Copyright 2022, Andrew Soltisz & Rengasayee Veeraraghavan, The Ohio State University. **Availability of data and material** Raw experimental data are backed up on Ohio State University servers and will be shared freely upon request. **Code availability** Example MATLAB code for SPACE, isotropic signal replacement, and synthetic image generation can be found on GitHub (https://github.com/andrewsoltisz/SPACE---Spatial-Pattern-Analysis-using-Closest-Events), and stand-alone code for SPACE can be found on MathWorks File Exchange (https://www.mathworks.com/matlabcentral/fileexchange/129464-space-spatial-pattern-analysis-using-closest-events). **Author Contributions** A. M. Soltisz: conceived of, developed, and implemented image analysis framework, and prepared the manuscript. Co-corresponding author. P. F. Craigmile: oversaw statistical validation of algorithm development and manuscript preparation. R. Veeraraghavan: oversaw the research project and assisted with algorithm development, and manuscript preparation. Co-corresponding author. **Ethics approvals** All animal procedures were approved by Institutional Animal Care and Use Committee at The Ohio State University and performed in accordance with the Guide for the Care and Use of Laboratory Animals published by the U.S. National Institutes of Health (NIH Publication No. 85-23, revised 2011).

## Abstract

The quantitative description of biological structures is a valuable yet difficult task in the life sciences. This is commonly accomplished by imaging samples using fluorescence microscopy and analyzing resulting images using Pearson’s correlation or Manders’ co-occurrence intensity-based colocalization paradigms. Though conceptually and computationally simple, these approaches are critically flawed due to their reliance on signal overlap, sensitivity to cursory signal qualities, and inability to differentiate true and incidental colocalization. Point pattern analysis provides a framework for quantitative characterization of spatial relationships between spatial patterns using the distances between observations rather than their overlap, thus overcoming these issues. Here we introduce an image analysis tool called *Spatial Pattern Analysis using Closest Events* (SPACE) that leverages nearest neighbor-based point pattern analysis to characterize the spatial relationship of fluorescence microscopy signals from image data. The utility of SPACE is demonstrated by assessing the spatial association between mRNA and cell nuclei from confocal images of cardiac myocytes. Additionally, we use synthetic and empirical images to characterize the sensitivity of SPACE to image segmentation parameters and cursory image qualities such as signal abundance and image resolution. Ultimately, SPACE delivers performance superior to traditional colocalization methods and offers a valuable addition to the microscopist’s toolbox.

## 1. Introduction

Structure-function relationships are fundamental to our understanding of biology. A notable case in point is the spatial distribution of biomolecules within a biological substrate, which is a key determinant of its biology and physiology. This perspective underpins much of our understanding of disease and its treatment, making the study of biological structures and their biomolecular patterns a common target of experimental characterization. Fluorescence microscopy is widely employed to observe and evaluate these patterns. The resulting images can then be analyzed using a variety of approaches, generally referred to as *colocalization*, which characterize biomolecule spatial distributions by their relative spatial association with some landmark, often another colabeled biomolecule. Recent decades have witnessed significant improvements in labeling technology and image quality, resolution, and acquisition rate, yet the accompanying spatial analyses have remained largely static despite their well-documented short comings.

Colocalization analyses are canonically categorized into either intensity-based or object-based methods. Intensity-based methods operate by comparing the brightness of individual pixels across an image’s spectral channels and are commonly performed using Manders’ co-occurrence or Pearson’s correlation paradigms (Bolte & Cordelières, 2006). Though popular and easy to implement, they are critically limited by their sensitivity to cursory differences in fluorescence signal attributes such as brightness and abundance (Wu et al., 2012). Additionally, they are only capable of assessing spatial relationships limited to direct signal overlap, making them resolution-dependent, incapable of describing spatial relationships between non-overlapping signals which may possess key spatial information about the biological structures under investigation, and inapplicable to monochromatic images, such as single channel fluorescence images or electron micrographs, which lack overlapping channels for pixel-wise comparisons.

Object-based analyses aim to overcome these limitations by utilizing the relative positions of observations detected via segmentation, called *objects*, to evaluate the spatial relationship of imaged signals based on either object-object overlap or inter-object distance thresholds (Bolte & Cordelières, 2006). These methods too are resolution-dependent and limited to analysis at only a single spatial scale in addition to being more difficult to implement and often, computationally expensive. None of these methods, intensity- or object-based, account for incidental colocalizations – those that occur simply by chance (Cordelières & Bolte, 2014).

The field of spatial statistics offers a variety of inter-object distance-based statistical frameworks, collectively called *Point Pattern Analysis*, for assessing spatial relationships that occur in excess of incidental colocalization (Cressie, 1993; Diggle, 2013). Here, we explore one such framework, a family of nearest neighbor (NN) distance distributions, as a means to quantitatively describe and compare spatial relationships from image-based microscopy data. We refer to our framework as *Spatial Pattern Analysis using Closest Events* (SPACE), and here, we illustrate its utility by comparison to traditional colocalization analyses, the Pearson’s and Manders’ paradigms, as well as application to experimental data of confocally imaged cardiomyocytes to measure the spatial relationship between their nuclei and select messenger ribonucleic acids (mRNA). Finally, we evaluate the sensitivity of this approach to cursory image signal characteristics and processing effects such as, segmentation parameters, object sample size, and signal concentration.

## 2. Materials and Methods

### 2.1. Sample Preparation and Imaging

Ventricular cardiomyocytes were enzymatically isolated from 8- to 12-week-old male C57BL6 mice and maintained in primary culture as previously described (Bogdanov et al., 2021). Individual mRNA molecules of interest were visualized with RNAScope Multiplex Fluorescent Reagent Kit v2 per manufacturer-recommended protocols. Briefly, isolated myocytes were fixed using paraformaldehyde, dehydrated in ethanol, and stored in 100% ethanol at -20 °C. Samples were then rehydrated for hybridization and incubation steps. Confocal imaging was performed using a Nikon A1R-HD laser scanning confocal microscope with individual fluorophores imaged sequentially to avoid spectral bleed-through.

### 2.2. Correcting Image Spatial-Anisotropy

We efficiently measure inter-event distances using distance transformations which require spatially isotropic images to produce accurate Euclidean distances. However, due to the elongation of the point-spread-function in the axial dimension, our confocal images have poorer resolution in the z-dimension. As a result, sampling is coarser in the axial dimension (z) than in the lateral dimensions (x, y), rendering the resulting digital images spatially anisotropic. While this could be corrected by simply resampling the image, the interpolation step will alter both signal morphology and the number of pixels that compose the signals, and thus reduce statistical power of subsequent analyses. Instead, we correct image spatial anisotropy, while avoiding these issues, using a technique we call *isotropic signal replacement* (see source code at (https://github.com/andrewsoltisz/SPACE---Spatial-Pattern-Analysis-using-Closest-Events).

First, the raw input image is segmented to identify signal-containing pixels. Then, a new blank image template is created with the same overall dimensions as the input image but with isotropic sampling, i.e. the z-dimension is sampled at the same spatial frequency as the lateral dimensions. Segmented pixels from the input image are then inserted back into this template image at pixel locations that are closest to the signal’s original Euclidean coordinates – in other words, pixels in the new image with centroid positions closest to those of the original signal-positive pixels in the raw image. Only these “replaced” pixels are used for subsequent analyses, thus maintaining the same sample size (number of signal-positive pixels) and signal morphology as the input image. It should be noted that positional rounding during the pixel replacement step can lead to small deviations in these pixel positions between the input image and the newly created image, thus altering measured distance. However, if the z-dimension is subdivided by a factor that is less than or equal to the lowest image resolution divided by root three, then the distance error between any two pixels within the new image will be less than the optical resolution of the image. Thus, any resulting errors in measured distances used for SPACE would fall below the least significant digit.

### 2.3. Image Segmentation

Signal masks from cardiomyocyte images were primarily generated using intensity thresholding (Xing & Yang, 2016). Specifically, cell body masks were composed of pixels whose brightness exceeded the mean by 10% or more. The cell body was identified as the largest resulting connected component which was then smoothed and filled using a morphological closing operation with a 5μm radius disk-shaped structural element. Nucleus masks were composed of pixels whose brightness exceeded the mean by at least five times the standard deviation. Any resulting connected components with volume less than 50 μm^3^ were removed and the remaining objects were individually smoothed and filled using a morphological closing operation with a 2μm radius disk-shaped structural element. mRNA masks were composed of pixels whose brightness was in the 98^th^ percentile if the image’s signal-to-noise ratio (SNR) was less than 40. Otherwise, pixels in the 99.6^th^ percentile were used. SNR was defined as the ratio of the 99.99% percentile brightness value to the 90% percentile value. Next, all connected components composed of fewer than 4-pixels were removed from the mask. Nucleus and mRNA mask pixels outside of their corresponding cell body mask were removed to restrict analysis to the intracellular space. All resulting masks were visually inspected to ensure accurate signal identification. Inaccurate masks were regenerated using manually derived brightness thresholds.

### 2.4. Empirical Distribution Functions

The x-coordinates of individual empirical cumulative distribution functions (CDF), which correspond to statistical G- and F-functions (Unwin, 1996), were defined as the unique distance values from the sampled distance transformations, sorted in ascending order. The y-coordinates were generated by first counting the number of events at each unique sorted distance then cumulatively summing the resulting counts before dividing each by the total event count. A count value of zero was included for a distance value of zero, if no events were found at this distance, to ensure continuity of the CDF over the full distance range. Likewise, a count of zero was included for any distance values where no observations occurred. The G-function, which is the CDF of observed nearest-neighbor distances for a specific signal component within an image, was generated by intersecting the mask for that signal component with the distance transformation of the landmark signal. The F-function, which is the CDF of nearest neighbor distances predicted under complete spatial randomness, was generated by intersecting the cell body mask with the distance transformation of the landmark signal. Individual delta CDFs were generated by first placing the paired G- and F-functions on a global x-coordinate scheme before subtracting F-function y-values from corresponding G-function values. The global x-coordinate scheme was defined as the union of both functions’ x-coordinates.

To summarize the central tendency of the distribution functions over individuals we calculated their median values at each distance. Specifically, individual CDFs were placed on a global x-coordinate scheme before computing the weighted median y-value at each distance. The global x-coordinate scheme was defined as the union of all individual functions’ x-coordinates and an event count of zero was assigned to each distance value that was not defined in the original individual CDFs before cumulatively summing event counts and normalizing to their total count. Median delta CDFs were calculated using the G- and F-functions defined over this same global x-coordinate scheme to ensure corresponding function values at the same x-coordinates.

### 2.5. Synthetic Image Generation

Synthetic images were created using a custom algorithm (see source code at (https://github.com/andrewsoltisz/SPACE---Spatial-Pattern-Analysis-using-Closest-Events) that generates two image-based point patterns, *X* and *Y*, such that the mean distance between each *Y*-event and its closest *X*-event can be statistically modulated using a single input parameter, *S* (**Supplemental Fig. 1**). Given a user specified image size, desired event count for each pattern, and target *Y→X* spatial proximity *S* within the range [-1,1], the *X*-pattern mask is first generated by assigning each *X*-event to its own unique and random pixel within the image, without replacement. Each *Y*-event is then assigned a distance-from-*X* using the inverse-CDF method and subsequently assigned, with replacement, to a random pixel within the image that satisfies its distance assignment. Here, each *Y*-event is assigned a random value within the range [-1,1] which is then used to interpolate the *Y→X* distance value from the CDF (**Supplemental Fig. 1D**) derived from the PDF (**Supplemental Fig. 1C**) of all distances composing the distance transformation of *X*’s mask which represents the distribution of *Y→X* distances expected under the CSR condition. The mean *Y→X* distance is tuned by using the spatial proximity parameter *S* as an input to a transfer function which subsequently biases the morphology of this assignment-PDF to select for smaller or larger distances (**Supplemental Fig. 1A**). When *S* is 0, the assignment-PDF is unaltered and *Y*-events will be assigned distances expected under CSR (**Supplemental Fig. 1B-E middle**). When *S* is negative, the assignment-PDF is transformed to increase the frequency of larger distances, thus *Y*-events will generally be assigned distances larger than expected under CSR (**Supplemental Fig. 1B-E top**). And when *S* is positive, the assignment-PDF is transformed to increase the frequency of smaller distances, thus *Y*-events will generally be assigned distances smaller than expected under CSR (**Supplemental Fig. 1B-E bottom**).

### 2.6. Pearson’s and Manders’ Colocalization Analyses

The Pearson’s correlation coefficient used for colocalization analysis was computed using one spectral channel as the x-variable and the other as the y-variable (Zinchuk & Grossenbacher-Zinchuk, 2014). This value was directly computed for the synthetically generated binary images, however, monochrome confocal images were first normalized to the maximum brightness of both images to reduce effects induced by differential fluorophore brightness between channels. Manders’ co-occurrence coefficient was calculated for synthetic binary images as the fraction of *Y*-pixels overlapped with *X*-pixels, and for experimental images as the fraction of segmented mRNA pixels overlapped with segmented nucleus pixels (Zinchuk & Grossenbacher-Zinchuk, 2014).

### 2.7. Statistics

A significance level of 0.05 was used for all statistical tests. The 2-sided Kolmogorov Smirnov (KS) test was used to statistically compare cumulative distribution functions. A weighted Student’s T-test was used for comparing samples of spatial association indices, with weights defined as the number of events composing the point pattern used to generate the underlying empirical distribution functions. To summarize the uncertainty, 5% and 95% quantiles were used for the estimated error envelopes of all distribution functions. The sample size for each experiment is indicated in their corresponding figure legends.

### 2.8. Code

All algorithms, including image segmentation, analysis, plotting, and generation of synthetic data were implemented in MATLAB (2023a. Natick, Massachusetts: The MathWorks Inc.). This code, as well as the raw data and their analysis results, will be provided upon request. Example code to generate synthetic images and to perform single-and multi-image SPACE analysis can be found on MATLAB File Exchange (https://www.mathworks.com/matlabcentral/fileexchange/129464-space-spatial-pattern-analysis-using-closest-events). Additional code for performing isotropic signal replacement and generating synthetic images can be found on GitHub ((https://github.com/andrewsoltisz/SPACE---Spatial-Pattern-Analysis-using-Closest-Events).

## 3. Results

### 3.1. Theory – Adapting Point Pattern Analysis for Image Data

We start by considering an image’s fluorescent signals *X* and *Y* as discrete spatial point patterns with their segmented signal pixels treated as discrete *events* contained within a finite *study area* defined by the image’s field of view or a region of interest (ROI) image mask for subregion analysis. Under this framework, we can use point pattern analysis, a powerful spatial statistics tool, to describe the spatial relationship between *X* and *Y* using distribution functions derived from their inter-event distances. Specifically, we leverage a theoretical bivariate point pattern analysis framework first described by Diggle & Cox, 1983. Using the point patterns *X* and *Y* whose event positions are defined by their segmentation masks (**Fig. 1A, top**), *X*’s spatial relationship to *Y* (*X→Y*), where *X* is the “subject” pattern and *Y* the “landmark” pattern, is described by *X*’s so-called bivariate G-function *G_X→Y_*(*r*), which defines the probability *P* of an arbitrary *X*-event existing within distance *r* from a *Y*-event. Adapting this for analysis of image data, we estimate *G_X→Y_* as the empirical CDF of distances between each pixel in *X*’s image mask and its closest pixel from *Y’s* mask. These distances can be efficiently calculated as the intersection of *X*’s mask with the distance transformation of *Y*’s mask (**Fig. 1A, middle**). Conveniently, the y-intercept of this function indicates the fraction of *X*-events that overlap with *Y*-events.

**Figure 1.**
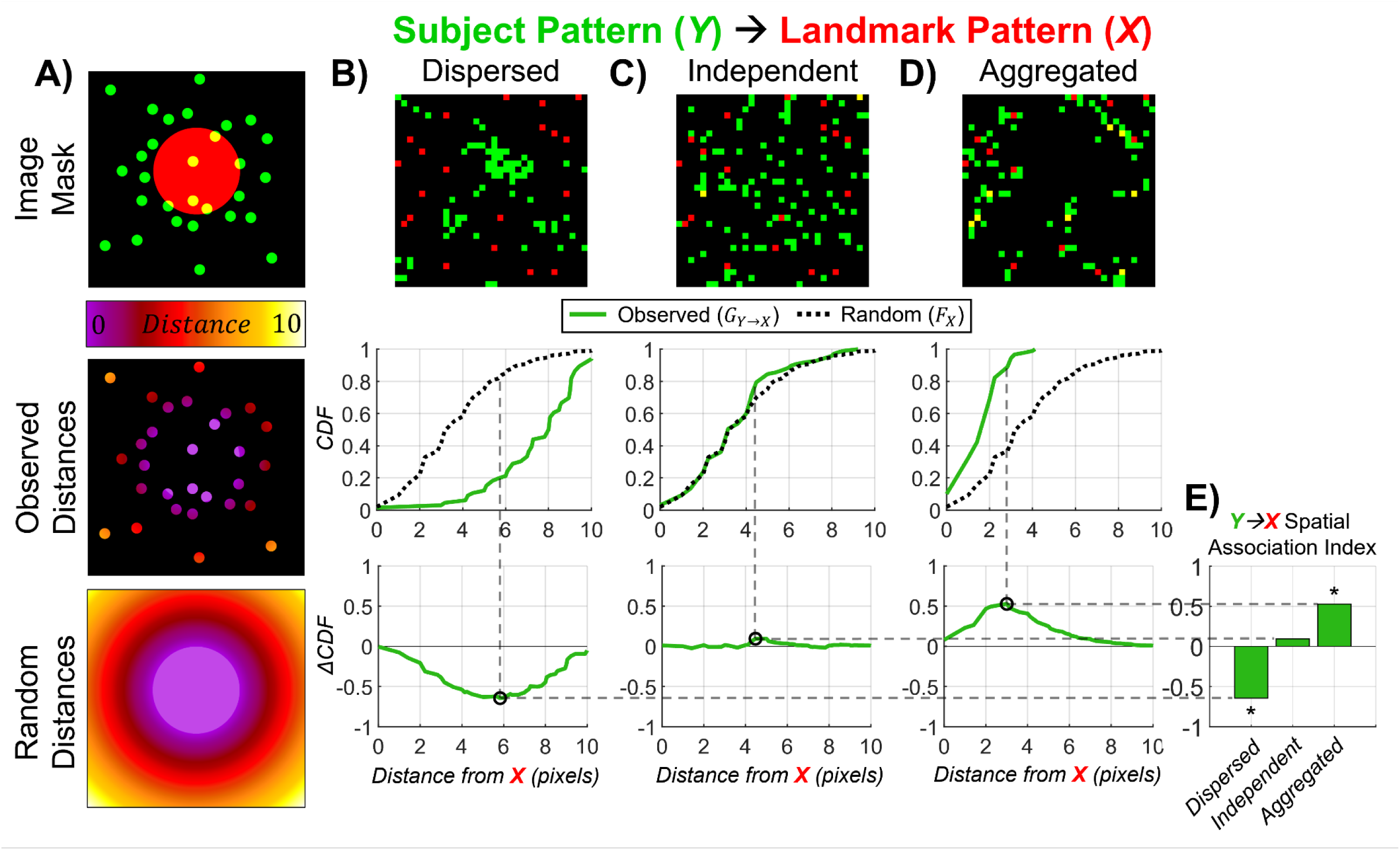
(**A top**) Cartoon image with two patterns, *X* (red) and *Y* (red). (**A middle**) Heatmap of observed distances between each *Y*-event and their nearest *X*-event. (**A bottom**) Heatmap of random distances that are the distance between each pixel in the image and their nearest *X*-event. (**B-D top**) 2D synthetic images exhibiting (**B**) dispersed, (**C**) independent, and (**D**) aggregated patterns. (**B-D middle**) Empirical CDFs of inter-event distances measured from the **B-D top** row images. Observed CDFs are plotted with solid green lines and random CDFs are plotted with dotted black lines. (**B-D bottom**) Delta CDFs representing deviation from CSR as a function of distances with black circles marking their peaks. (**E**) Resulting spatial association indices with asterisks indicating significant deviation from CSR evaluated using a 2-sided KS test. Horizontal and vertical dashed gray lines respectively indicate the values and distances of delta function peaks from which spatial association indices were derived.

Scalar summary statistics that describe *X→Y* could be derived directly from *G_X→Y_*, however they will be influenced by cursory image characteristics that affect inter-event distances such as the event concentration of both patterns and the geometry of the study area. To describe spatial relationships independent of these other factors, *G_X→Y_* can be compared to the G-function *Ĝ_X→Y_* that would be produced, given the specific state of these cursory factors, if pattern *X* were indeed the result of a homogenous Poisson point process, an idealized point pattern which represents complete spatial randomness (CSR) between *X* and *Y*. Under this condition, *X*-event locations are independent samples from the uniform random distribution and the number of events contained by an arbitrary circular region with radius *r* is a Poisson distribution. Deviation of process *X*’s original “observed” G-function *G_X→Y_* from its “random” counterpart *Ĝ_X→Y_* occurs when *X→Y* is not CSR, but instead *X* is aggregated or dispersed relative to *Y*. It will therefore be useful to compare these distributions as a means to describe the statistical spatial relationship between fluorescence signals *X* and *Y* independent of cursory image qualities.

*Ĝ_X→Y_* could be derived analytically, though it may be difficult, if not impossible, to define an equation that accounts for effects induced by study areas with complex geometries, a common occurrence in biology, such as when the study area is defined by the irregular and tortuous surface of a cellular membrane. *Ĝ_X→Y_* could also be approximated using Monte Carlo-style simulations, though this approach becomes computationally inefficient when applied to large or finely discretized study areas such as in images produced from high-resolution microscopy. An alternative CSR fiducial distribution is process *Y*’s so-called univariate F-function *F_Y_*(*r*), which describes the probability of an arbitrary point in space existing within distance *r* from a *Y*-event (Cressie, 1993; Diggle, 2013; Dixon, 2001). Adapting this for analysis of image data, we derive an estimate of *F_Y_* as the eCDF of distances between each pixel from the ROI mask and its closest pixel from *Y’s* mask. These distances, and therefore the F-function, can be efficiently calculated as the intersection of the ROI mask with the distance transformation of *Y*’s mask (**Fig. 1A, bottom**). Conveniently, the y-intercept of this function indicates the expected fraction of *X*-events that would overlap with *Y*-events under the CSR condition. Furthermore, when *X*-event positions are independent of *Y* and their inter-event NN distances match those expected under CSR, both *G_X→Y_* and *F_Y_* will converge to *Ĝ_X→Y_*, meaning *G_X→Y_* and *F_Y_* will be equal and their delta function, produced by subtracting *F_Y_* from *G_X→Y_*, will be a horizontal line along the x-axis (**Fig. 1C**). When *X* is dispersed from *Y* and their NN distances are greater than expected under CSR, *G_X→Y_* will be right shifted from *F_Y_* and their delta function will exhibit a prominent negative peak (**Fig. 1B**). And when *X* is aggregated with *Y* and their NN distances are less than expected under CSR, *G_X→Y_* will be left shifted from *F_Y_* and their delta function will exhibit a prominent positive peak (**Fig. 1D**). These two distributions can be rigorously distinguished using a 2-sided KS test and the statistical spatial association index of *X→Y* parameterized as the peak value of their delta function, where dispersion produces negative values, aggregation positive values, and CSR values at or near zero (**Fig. 1E**).

Pattern *Y* can also exhibit its own unique spatial relationship with *X* (*Y→X*) which can be evaluated by performing this analysis from the perspective of *Y* instead of *X* (**Supplemental Fig. 2**). Here, *Y*’s “Observed” G-function *G_Y→X_* is produced using the distances between each pixel in *Y*’s image mask and its closest pixel from *X*’s mask, and the fiducial “random” F-function *F_X_* is produced using the distances between each pixel from the ROI mask and its closest pixel from *X*’s mask. Both perspectives must be analyzed to fully describe the bidirectional spatial relationship between *X* and *Y* (*X*↔*Y*), though in certain cases, only one perspective may be informativeor relevant, such as when one pattern, let’s say *Y,* is monolithic and the other is composed of many events throughout the study area. In this case, the *Y→X* analysis will be severely limited by the lack of positional diversity of *Y*-events, thus causing many *X*-events to not qualify as nearest neighbors to *Y* so the analysis will not capture their likely valuable spatial information. Conversely, the *Y*-events ignored in the corresponding *X→Y* analysis will still be in close proximity to those *Y*-events that do qualify as NN to *X*-events, thus their positional information will still largely be captured.

### 3.2. Extension to Multiple Images

Microscopy studies often utilize many replicate images to statistically describe the structural behavior of populations, making it vital for downstream image analyses to be capable of collating and statistically comparing groups of images. We have therefore extended this spatial analysis framework to do just that (**Fig. 2, Supplemental Fig. 3, 4**). Given a set of images sampled from a population (**Fig. 2A**), the central tendency of their distribution functions is estimated by the weighted median of the individual functions and the error estimated by the weighted upper and lower quantile envelope (**Fig. 2B-C**). Specifically, a function’s weighted median is defined as the weighted 50% quantile of function values (y-coordinates) at each distance (x-coordinate), where at each distance, the individual function values are first sorted in ascending order, then the quantile function value is chosen as the first ordered function value to cause the cumulative sum of individual function weights to exceed the specified quantile. Here, a function’s weight is calculated as its point pattern event count divided by the sum total event count across all functions in the set. Additionally, the upper and lower quantiles can be chosen according to the required level of confidence; 95% upper and 5% lower quantiles were used in this study. This approach is defined for both the observed and random distributions, as well as their delta function.

**Figure 2.**
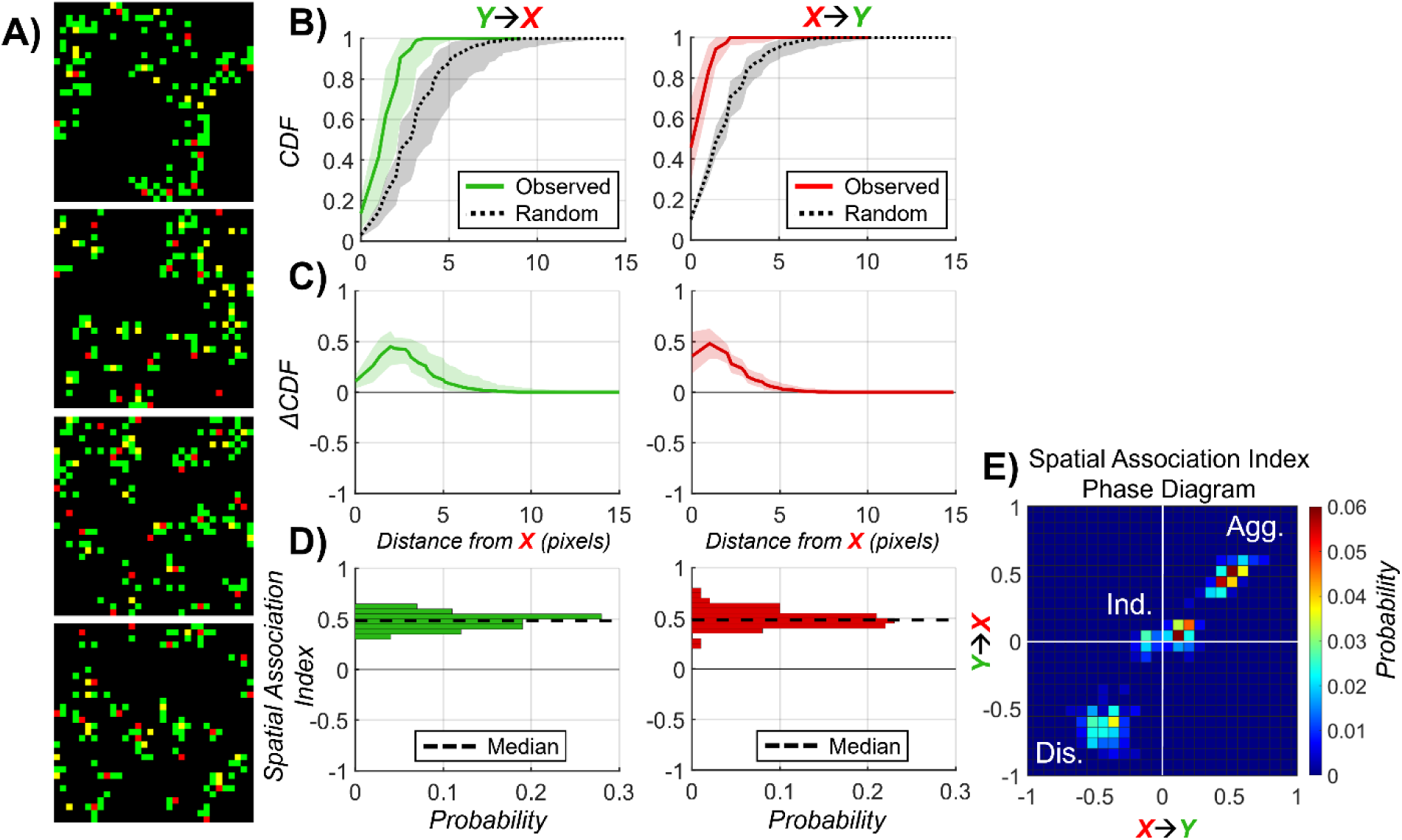
(**A**) Representative 2D synthetic images exhibiting aggregation between pattern-*X* (red) and pattern-*Y* (green) patterns. (**B**) Weighted median CDFs and (**C**) delta CDFs for both perspectives of SPACE analysis. Shaded regions indicate upper 95% and lower 5% weighted quantiles. (**B**) Observed CDFs are plotted with solid lines and random CDFs with dashed lines. (**D**) Histogram of *Y→X* (green) and *X→Y* (red) spatial association indices measured from the entire batch of synthetic images with spatially aggregated patterns. Dashed black lines indicate distribution medians. (**E**) Spatial association index phase diagram plotted as the bivariate histogram of *X→Y* spatial association indices for each image on the x-axis and *Y→X* spatial association indices on the y-axis. Images with dispersed, independent, and aggregated patterns form significantly different clusters indicated by Bonferroni-corrected pairwise weighted Student’s T-tests. Each group was composed of 100 images.

The spatial relationship between two point patterns sampled by a set of images can then be globally evaluated for CSR using their delta function envelope. If the envelope ever positively exceeds zero, then there is sufficient evidence to conclude that the relationship is aggregated (**Fig. 2C**). If there is a negative escape from zero, the relationship is dispersed (**Supplemental Fig. 4C**). Lastly, if the envelope overlaps with zero over the entire distance range, then the relationship is considered CSR (**Supplemental Fig. 4G**). Likewise, the spatial relationship can be statistically differentiated between two experimental groups of images by comparing their samples of individual spatial association values with a weighted Student’s t-test, using the aforementioned definition for sample weights (**Fig. 2D**). Finally, the bidirectionality of the spatial relationship exhibited by these two patterns can more easily be visualized using a type of phase diagram, where the *X→Y* spatial association index for each image is plotted on the x-axis and their corresponding *Y→X* spatial association index on the y-axis (**Fig. 2E**). Here, mutually aggregated relationships cluster in the upper right quadrant, mutually dispersed in the lower left quadrant, and mutually CSR relationships cluster around the origin.

### 3.3. Example Application –Distribution of mRNAs within Cardiomyocytes

Canonically, membrane proteins are thought to be synthesized via the secretory protein trafficking (SPT) pathway, where nascent mRNA are translated in the peri-nuclear rough endoplasmic reticulum (ER) and resulting proteins are processed and transported from the Golgi network to their sites of membrane insertion (He et al., 2020). However, recent studies of neurons, a large and highly differentiated cell type, demonstrate the presence of local membrane protein synthesis, where mRNA molecules themselves are trafficked from the nucleus to distinct cellular locations such as axon end segments and retained locally for on-demand translation (Holt et al., 2019). This alternative production pathway allows for faster tuning of protein synthesis in response to local signaling cues, like neuronal growth cone motility, which would otherwise be inefficient, if not unfeasible, using the slow kinetics of nuclear signaling and protein trafficking along axons that may extend beyond a meter in length (Dalla Costa et al., 2021). Given cardiomyocyte’s similarly large size, densely packed cytosol, and vital importance of membrane protein spatial distributions (Bentzinger et al., 2012), we hypothesized that they too synthesize membrane proteins in a similarly distributed fashion. Here, we demonstrate SPACE’s ability to address this practical biological question by measuring cardiac membrane protein mRNA’s spatial association with cell nuclei using our previously published experimental data (Bogdanov et al., 2021).

Cell nuclei and mRNA encoding for key cardiac cell membrane proteins (Cx43 protein encoded by Gja1 mRNA, Na_V_1.5 encoded by Scn5a, Ca_V_1.2 encoded by Cacna1c) and sarcoplasmic reticulum membrane proteins (SERC2a encoded by Atp2a2, RyR2 encoded by Ryr2) were fluorescently labeled in murine cardiomyocytes (**Fig. 3A**). Following image segmentation, the *mRNA→Nucleus* spatial relationship for each cell was measured using SPACE, with mRNA masks used as the “subject” pattern and nuclei masks as the “landmark” pattern. The cell body mask, derived from the nuclear fluorescence signal, served as the ROI which defined the space used to derive the random distribution function *F_Nucleus_*. SPACE analysis revealed all mRNA species distributed throughout the entire cell, although to varying degrees (**Fig. 3B**), illustrating clear deviations of membrane protein mRNA translation away from the canonical peri-nuclear SPT pathway. The resulting spatial association indices derived from comparing observed and random distribution functions further revealed mRNA species stratifying into varying nuclei-centric distributions, with Atp2a2 being CSR relative to the nucleus, Ryr2 exhibiting slight nuclear aggregation, Scn5a and Cacna1c with intermediate levels of nuclear aggregation, and Gja1 with near-maximal nuclear aggregation (**Fig. 3C-D**). Repeating this analysis from the perspective of the nucleus revealed a mutual bidirectional *mRNA*↔*Nucleus* spatial relationship, with the spatial association index phase diagram showing a high degree of alignment with line of identity (**Supplemental Fig. 5**).

**Figure 3.**
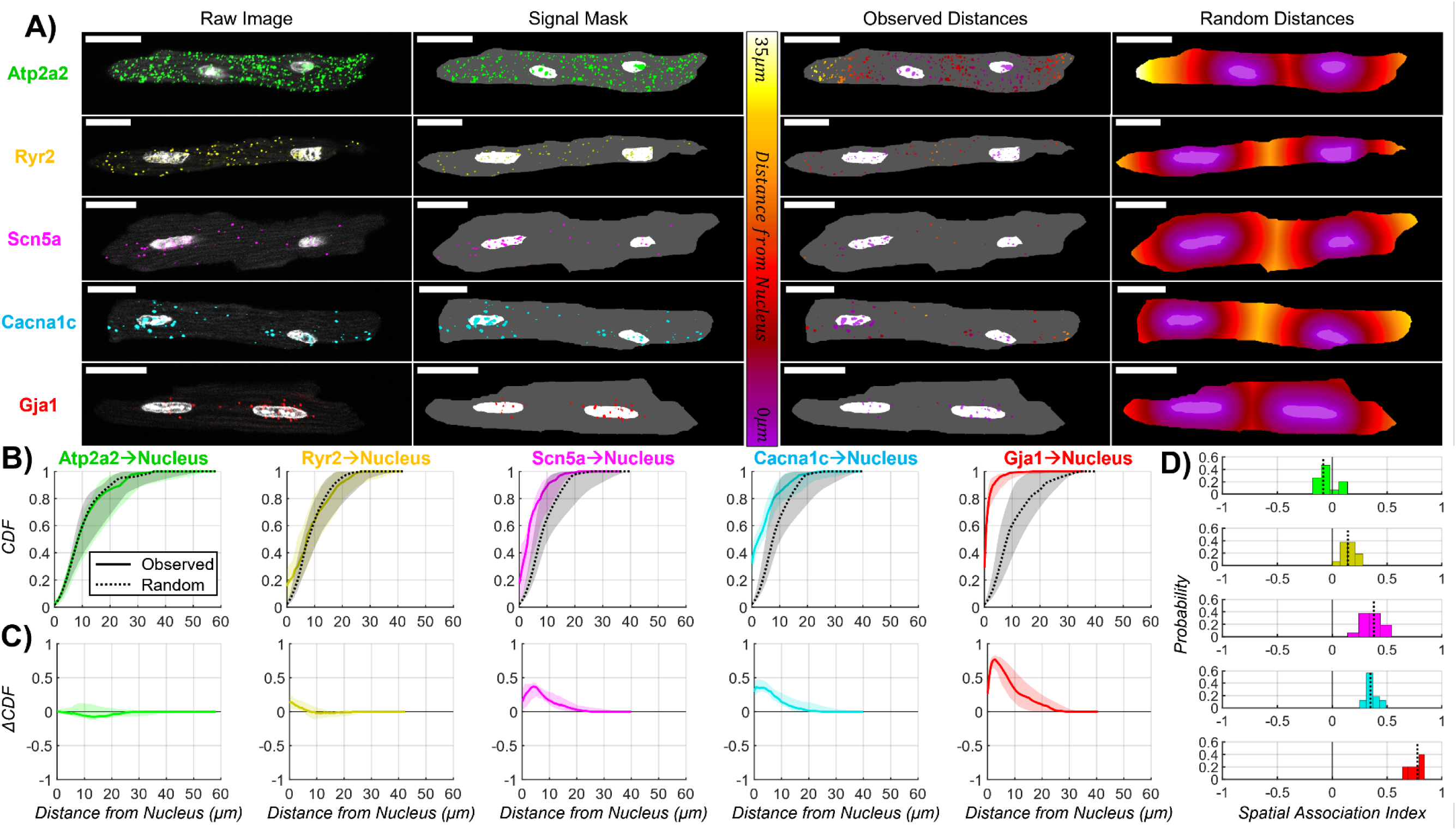
(**A left**) Representative 2D planes from 3D confocal images of individual cardiomyocytes with fluorescently labeled nuclei (white) and mRNAs Atp2a2 (green), Ryr2 (yellow), Scn5a (magenta), Cacna1c (cyan), and Gja1 (red). Scale bars are 20μm. (**A middle-left**) Image masks indicating event pixels for mRNA (colors) and nuclei (white) and the cell body as the study area (grey). (**A middle-right**) heatmaps of observed distances between each mRNA mask pixel and their closest nucleus pixel. (**A right**) heatmaps of random distances between each pixel in the study area and their closest nucleus pixel. (**B**) Weighted median CDFs and (**C**) delta CDFs for the *mRNA→Nucleus* SPACE analysis. Shaded regions indicate upper 95% and lower 5% weighted quantiles. (**B**) Observed CDFs are plotted with solid lines and random CDFs with dashed lines. (**D**) Histograms of *mRNA→Nucleus* spatial association indices for individual images with dashed black lines indicating medians. *mRNA→Nucleus* spatial association indices were compared between mRNAs using Bonferroni-corrected pairwise weighted Student’s T-tests (**Supplemental Table 2**). (Gja1: n = 10 cells from 3 hearts; Scn5a: n = 16 cells from 3 hearts; Cacna1c: n = 16 cells from 3 hearts; Ryr2: n = 16 cells from 3 hearts; Atp2a2: n = 15 cells from 3 hearts)

### 3.4. Validation and Sensitivity Analysis

The utility of SPACE is dependent on its ability to produce accurate estimates of the observed and random distribution functions. Generally, the more events captured within the ROI, the better the function estimates will be. We therefore characterized the stability of SPACE’s output spatial association index to changes solely in event sample size by randomly thinning the mRNA signal from individual cells labeled for the two mRNA species with the most extreme nuclear associations: Atp2a2 with the lowest spatial association index and Gja1 with the highest (**Fig. 4A-C, Supplemental Fig. 6A-C**). Before measuring distances, randomly selected pixels were deleted from the mRNA mask. This was repeated ten times for each thinning level which ranged between 0% and 99% deletion of the original signal. Although the variability of measured association increased as mRNA event sample size was reduced (**Fig. 4B**), the central tendency of the measurement remained stable even over extreme thinning regimes, with the mean absolute percent error in nuclear association remaining lower than 0.05% of the metric’s dynamic range for all thinning levels less than 80% (**Fig. 4C**). Thus, SPACE results are robust to a wide range of sample sizes.

**Figure 4.**
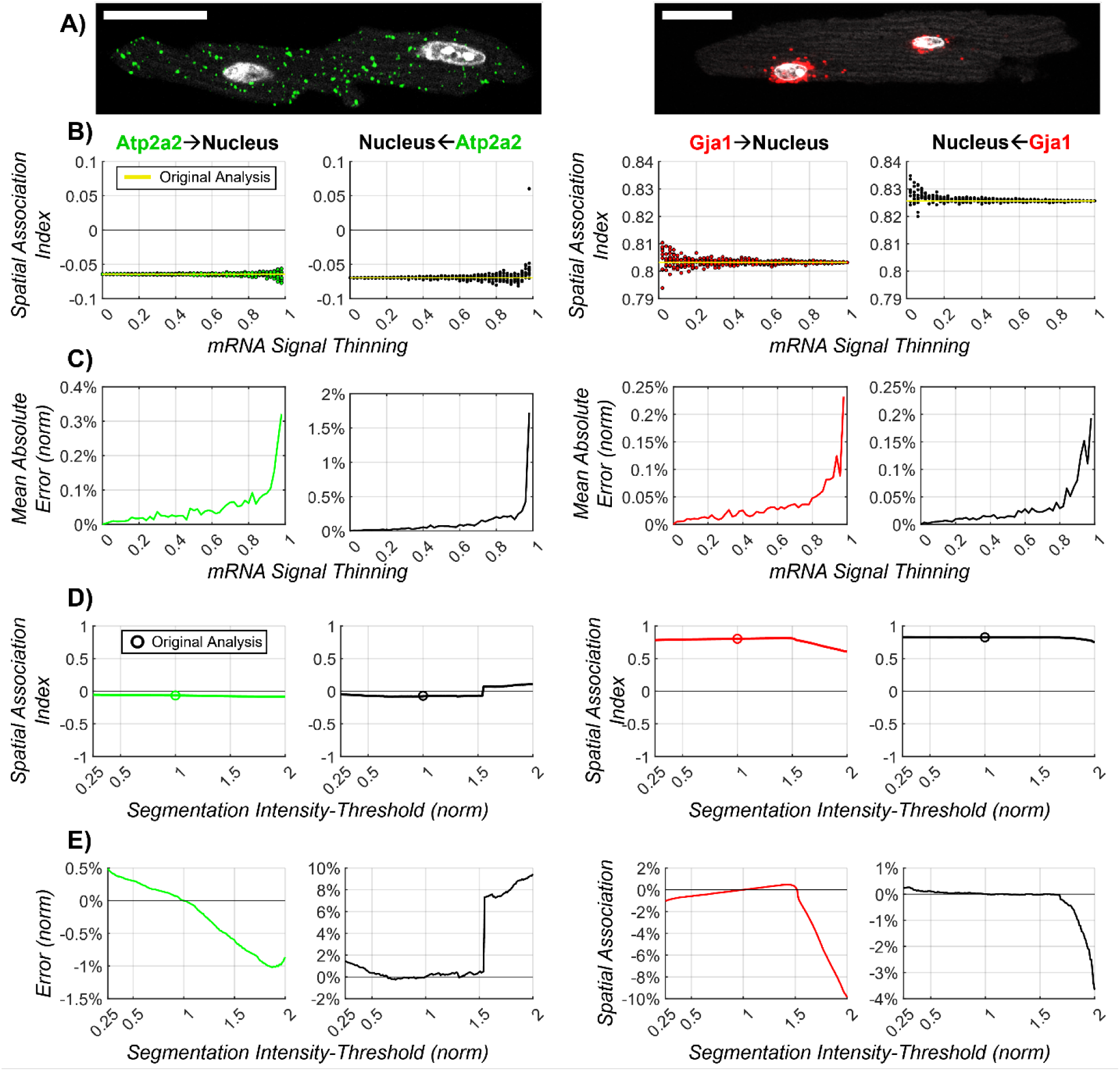
(**A**) 2D planes from 3D confocal images of individual cardiomyocytes with fluorescently labeled nuclei (white), Atp2a2 mRNA (green), and Gja1 mRNA (red). Scale bars are 20μm. (**B**) Scatter plots of *mRNA→Nucleus* (green or red) and *Nucleus→mNRA* (black) spatial association indices as a function of the level of mRNA signal thinning. Random signal thinning was varied between 1% and 100% with 1% increments and repeated 10 times per level. Yellow lines indicate the original spatial association index. (**C**) Mean absolute error of spatial association indices for the 10 repeats and normalized to index’s dynamic range. (**D**) Spatial association index as a function of mRNA segmentation intensity threshold. The threshold was varied between 25% and 200% with 1% increments. Circles indicate the original spatial association index. (**E**) Difference between threshold-adjusted and original spatial association index normalized to the index’s dynamic range.

Brightness thresholding-based image segmentation was the primary method employed to identify point pattern pixels from the images of mRNA-labeled cardiomyocytes. As with any explicit segmentation approach, parameter values, or the functions used to derive them, are manually chosen to achieve the best representation of the underlying signal, as subjectively assessed by the user. To evaluate the extent to which SPACE results are affected by the brightness thresholds we used for segmentation, we performed a segmentation sensitivity analysis on the same two Atp2a2 and Gja1 labeled cells (**Fig. 4D-E, Supplemental Fig. 6D,E**). The brightness threshold was varied between 25% and 200% of the original threshold and the effect on *mRNA→Nucleus* spatial association measured. Spatial association indices remained stable over a wide range of brightness thresholds (**Fig. 4D**), with the error remaining below 1% of the dynamic range between thresholds 50% to 150% of the original value (**Fig. 4E**). Thus, there is a wide range of brightness thresholds for this data that will produce nearly identical SPACE results.

Next, we sought to verify the independence of spatial association indices to point pattern concentrations, as inter-event distances will generally decrease with increasing concentration. For each cell labeled for the mRNAs with the largest variability in concentration, Atp2a2 and Gja1, we performed linear regression on the measured *mRNA→Nucleus* spatial association indices as a function of mRNA signal concentration (**Fig. 5A**). Here, concentration was defined as the ratio of mRNA pixel count to ROI pixel count. The slope of this line was not significantly different than 0, indicating no dependency between spatial association and concentration for these data (**Fig. 5B**). We also compared SPACE performance with the most commonly used spatial analyses in biological microscopy, Pearson’s and Manders’ colocalization. Manders’ measure of co-occurrence did not significantly vary with mRNA concentration, however Pearson’s measure of spatial correlation was indeed significantly affected by concentration (**Fig. 5**). This study suggests that SPACE and Manders’ are superior spatial analyses for this experimental data set while Pearson’s was noticeably affected by cursory signal characteristics, thus degrading its utility as an objective measure of spatial proximity.

**Figure 5.**
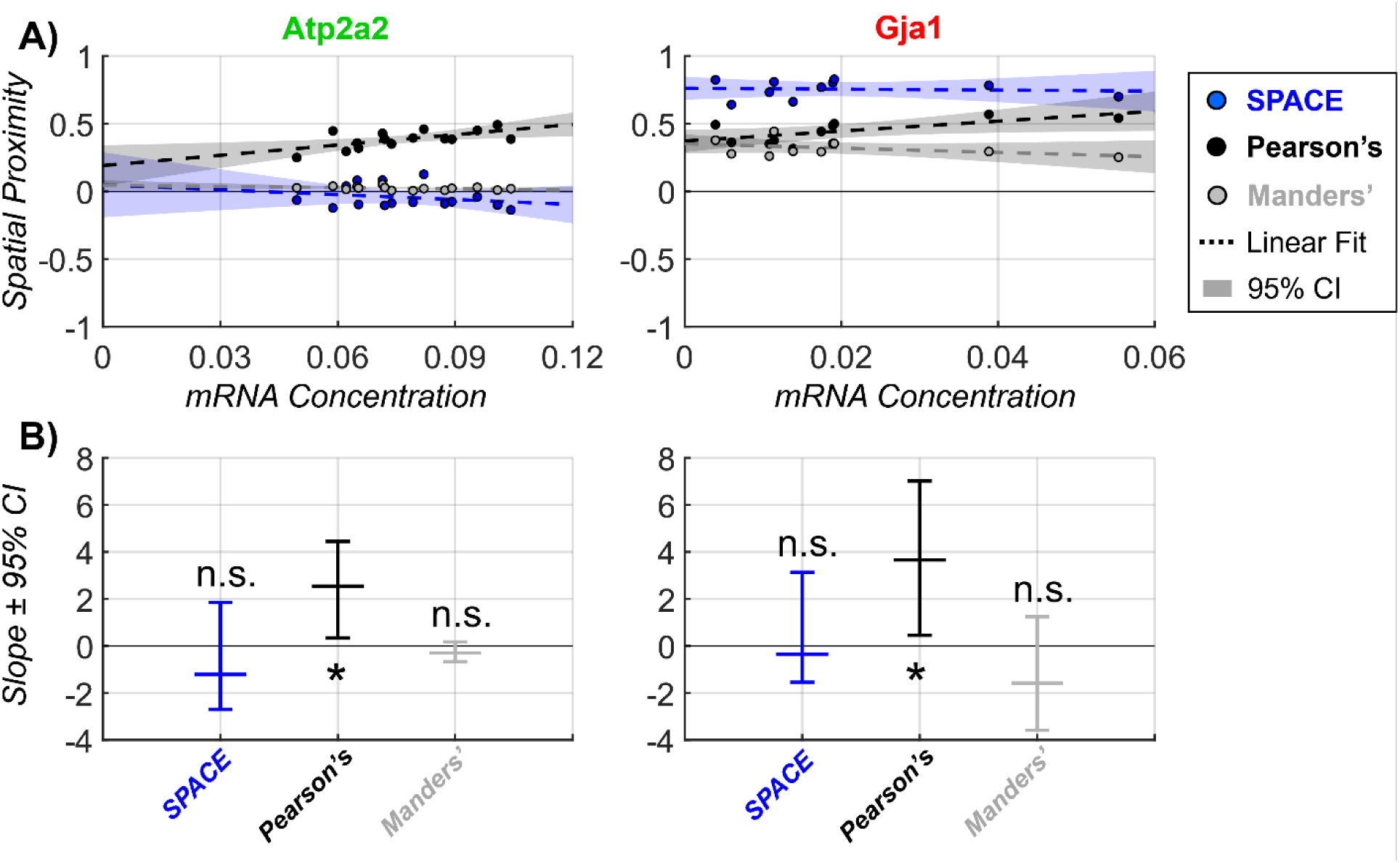
(**A**) Scatter plot of measured *mRNA→Nucleus* spatial proximity as a function of mRNA concentration for individual cardiomyocytes labeled for Atp2a2 and Gja1 mRNA. SPACE’s spatial association index is marked by blue, Pearson’s correlation is black, and Manders; co-occurrence is gray. Linear regression lines are marked by dashed lines and their 95% confidence interval is shaded. (**B**) Plots of the linear regression slope with error bars representing the slope’s 95% confidence interval. Asterisks indication a slope that is significantly different than zero and n.s. is not significantly different than 0.

However, mRNA concentration and its spatial relationship with the nucleus may be biologically correlated, so we also performed a similar study using synthetically generated images to control for potential confounding factors (**Fig. 6**). Within a 100x100-pixel image, events composing the point pattern *X*, represented by red pixels, were randomly seeded into the image before *Y*-events, represented by green pixels. The latter were assigned to specific pixels using a custom inverse CDF-based method that tunes the mean distance between *Y*- and *X*-events using a single spatial proximity parameter *S*. With *S* held at 0.8 to produce a mean Y*→*X distance less than CSR, and the concentration of pattern-*Y* held at 10%, the concentration of pattern-*X* was varied between 1% and 100% (**Supplemental Movie 1**), and *Y→X* spatial proximity measured with SPACE, Pearson’s, and Manders’ analyses (**Fig. 6A**). SPACE was the most stable measure of spatial proximity, as measured by its variance over the entire concentration range normalized to the metric’s dynamic range (**Fig. 6B**). Additionally, we assessed the sensitivity of each analysis method to changes in *Y→X* mean distance. With *X*- and *Y*-pattern concentrations both held at 10%, S was varied between -1 and 1 (**Supplemental Movie 2**) and *Y→X* spatial proximity measured with SPACE, Pearson’s, and Manders’ analyses (**Fig. 6C**). SPACE was the most sensitive measure of spatial proximity to changes in mean inter-event distance, with the largest variance over the entire range of *S* (**Fig. 6D**). Ultimately, SPACE was least effected by cursory image qualities and most sensitive to changes in their spatial relationships, suggesting it is a superior method for spatial analysis of image-based data.

**Figure 6.**
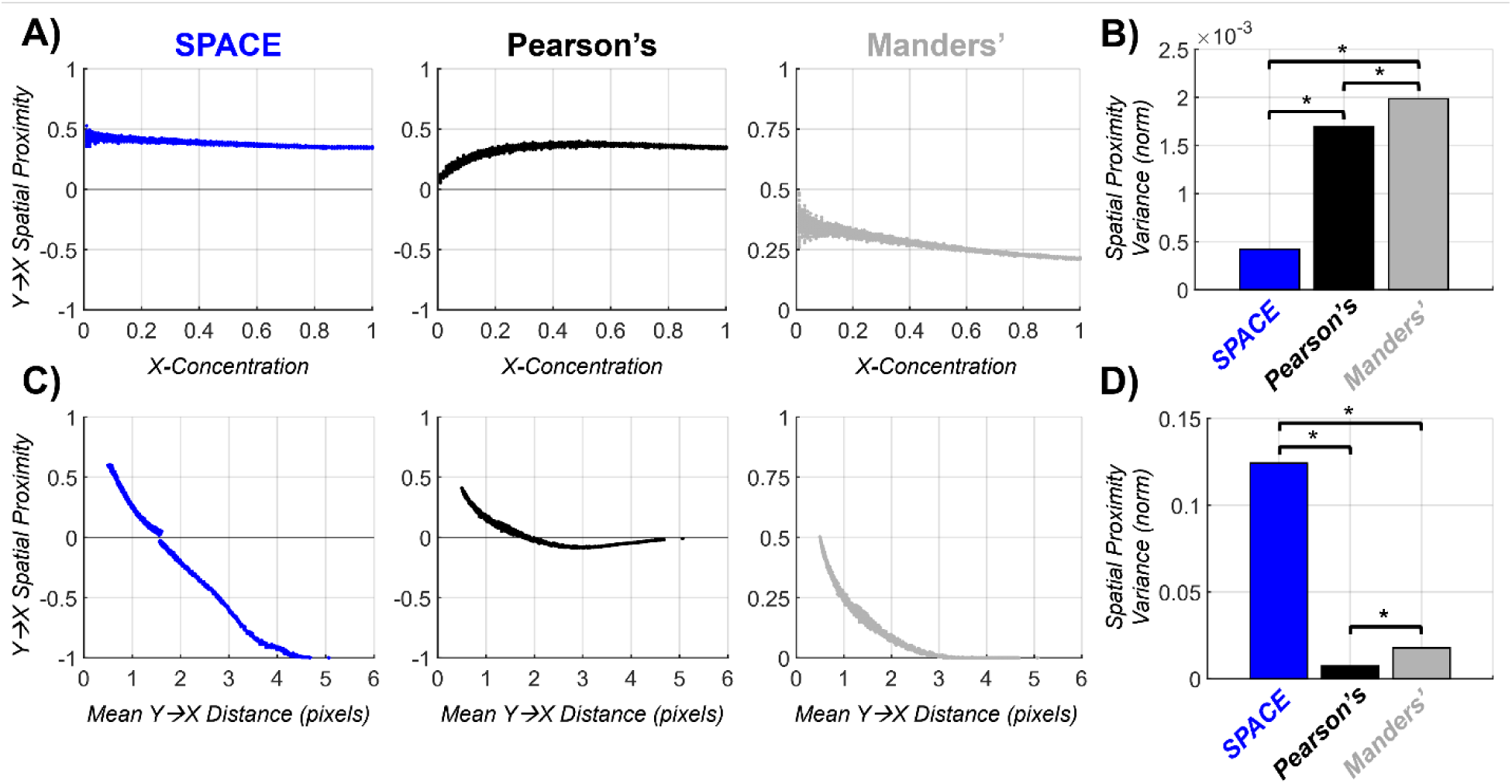
(**A**) Measured spatial proximity for synthetic pattern-Y relative to pattern-*X* as a function of *X*-concentration for individual synthetic images. *X*-concentration was varied between 1% and 100% by increments of 1%. 50 unique synthetic images were generated per concentration level. SPACE’s spatial association index is marked by blue, Pearson’s correlation is black, and Manders; co-occurrence is gray. (**B**) Variance of measured spatial proximity over the entire range of X-concentrations, normalized to the dynamic range of each spatial proximity metric. (**C**) Measured *Y→X* spatial proximity as a function of the mean *Y→X* nearest neighbor distance for synthetic images whose input *Y→X* spatial proximity was varied between -1 and 0.995 by increments of 0.005. 10 unique synthetic images were generated per spatial proximity level. (**D**) Variance of measured spatial proximity over the entire range of input *Y→X* spatial proximity, normalized to the dynamic range of each spatial proximity metric. Asterisks mark statistically significant differences in variance indicated by Bonferroni-corrected pairwise 2-sided F-tests.

## 4. Discussion

The analysis of fluorescence microscopy images to discern spatial patterns within imaged signal components relative to each other is a fundamental tool in the biologist’s arsenal. As discussed above, the long-used conventional colocalization analyses provide limited insight, while also being dependent on cursory differences in fluorescence signal attributes such as brightness and abundance (Wu et al., 2012). Further, they are resolution-dependent, losing meaning as the resolution of light microscopy techniques extends below the size of biomolecules. To address these issues, we introduce here *Spatial Pattern Analysis using Closest Events* (SPACE), a novel image analysis framework, which adapts a paradigm of point pattern analysis to quantitatively measure the degree of non-random spatial association exhibited between two sets of signal components within image data. In this study, we describe its implementation, validate is performance characteristics, verify its robustness with respect to cursory signal properties, and demonstrate its utility by analyzing the spatial association between cardiomyocyte mRNA and nuclei. We provide evidence that our novel technique is superior to the two most commonly used spatial analyses in biological microscopy studies, Pearson’s and Manders’ colocalization.

SPACE offers the biologist several key advantages by virtue of its design and implementation. Most notably, the point process analysis approach employed by SPACE provides rich and robust quantification of spatial patterns occurring over all length scales contained within the image. In contrast, binary analyses based on super-position of signals, such as Pearson’s and Manders’ colocalization, capture very limited information while being resolution-dependent and limited to analysis at only a single spatial scale. While characterizing spatial relationships using inter-object distances alleviates these problems, such methods require a robust statistical framework to ensure rigorous and informative analysis. Simply defining an *a priori* distance threshold, under which objects are considered colocalized, is undesirable, as it requires the user to choose what they deem to be a functionally relevant distance, thus requiring prior knowledge of the system under study. This information is often unavailable or unknown, thus the chosen threshold will likely introduce bias into the analysis and again limit assessment of spatial relationships to this one spatial scale. By being agnostic to specific length scales, SPACE offers a convenient solution to this problem. Furthermore, the G- and F-functions calculated by SPACE can be further probed with computational models to uncover the mechanistic spatial processes which drive the observed distributions, as we have previous demonstrated (Bogdanov et al., 2021).

Through its use of distance transformations for calculating nearest neighbor distances, SPACE avoids the need for computationally expensive Monte Carlo-style simulations to estimate the CSR distribution function, thus rendering the approach suitable for large datasets as well as real-time analysis in conjunction with image acquisition. This holds important implications, given recent advances in smart microscopy and automated, adaptive image acquisition strategies (Barentine et al., 2023; Daetwyler & Fiolka, 2023). Additionally, SPACE is designed to facilitate application to multi-sample datasets and inter-group comparisons, making for robust, repeatable results.

In addition to the aforementioned advantages over conventional colocalization methods, SPACE is specifically designed to overcome specific issues that confound results obtained by the latter. We demonstrate how two cursory, and often highly variable, signal qualities, signal concentration and sample size, corrupt other analyses but do not significantly affect outputs from SPACE. Additionally, SPACE requires little user input due to its non-parametric nature, although signal segmentation is left to the user’s discretion. Equally, the modular nature of SPACE allows users to replace the simple intensity-based thresholding approach used for segmentation with more sophisticated methods selected to match the characteristics of specific applications.

In contrast to traditional colocalization analyses, SPACE is not restricted to analysis of only multi-channel fluorescence microscopy images. Its use of inter-event distance allows for the characterization of spatial relationships between non-overlapping signals, thus making it amenable to assessment of any digital image with arbitrarily many spectral channels, whether it be a micrograph captured by a microscope or an image taken with a personal smartphone. So long as image components can be segmented into masks, their spatial relationship can be characterized using SPACE, although anisotropic or distorted images will need to be spatially calibrated and corrected to ensure accurate measures of distance. Furthermore, this style of analysis can even be applied to non-image data, such as pointillist data of fluorophore coordinates derived from single-molecule localization microscopy, or even generalized for application to non-spatial data, where events can instead be thought of as samples whose positions are defined within an n-dimensional variable space rather than by Euclidean coordinates, though an appropriate distance metric will need to be defined. In either of these cases, the analysis will likely carry on increased computational load as the brute-force and graph-based methods for identifying nearest neighbors from continuous data have much larger time complexities compare to the modern linearly-timed algorithm used for calculating distance transformations of discrete image data (Maurer et al., 2003).

While SPACE offers the user multiple advantages as discussed above, there are important factors to be considered to ensure robust results. Foremost is ensuring input data are of good quality and relevant to the research question(s) at hand. The user must take care to verify fidelity of fluorescence signals to the biological structure(s)/process under study. Related to this, images must be collected so that the field of view captures a single pattern, where possible. If not, it is important to bear in mind while interpreting results that they reflect information aggregated from all captured spatial relationships. Although we demonstrate here that SPACE is insensitive to fairly large changes in the intensity threshold used for segmentation, this should be directly verified for use cases where the properties of the input data and/or segmentation are significantly different. Segmentation methods should be employed to select pixels that are deemed as appropriate markers for their signals of interest, while avoiding all other pixels, as much as possible. To this end, validating segmentation strategies using known ground truth, such as synthetic data, and performing parameter sensitivity analyses are sound practices. In this context, it should be noted that such validation is often not possible for machine learning-based black box segmentation techniques. Thus, caution is advised with their use as it is difficult to verify the fidelity of segmentation results. Last but not least, CSR is almost certainly a biologically unachievable spatial relationship, but using it as a benchmark serves as a good first step towards assessing whether there are non-random spatial heuristic(s) dictating the relative positions of imaged signals. This null hypothesis could be substituted for a more complicated distribution chosen based on prior knowledge of the system under study. However, other NN distance distributions tend to be much harder to generate, both analytically and computationally (Dixon, 2001).

## 5. Conclusion

In summary, we present here a novel pipeline, called *Spatial Pattern Analysis using Closest Events* (SPACE), that offers significant improvements over conventional spatial analyses due to its quantitative nature, stability, and computational efficiency. This approach is also agnostic to the imaging modality or resolution achieved. Thus, its applicability is limited to neither fluorescence images nor the microscopic spatial scales.

## Acknowledgements

The authors wish to thank Mr. Vladimir Bogdanov, Dr. Jonathan P. Davis, and Dr. Sandor Gyorke for providing isolated murine cardiac myocytes for imaging studies and Dr. Raghu Machiraju and Dr. Seth Weinberg for valuable discussions, which motivated the development of *Spatial Pattern Analysis using Closest Events*.

---

Supplemental Video 1.mp4

**Supplemental movie 1**. Video showing representative synthetic images for each *X*-concentration from main figure 6 panels A and B.

Supplemental Video 2.mp4

**Supplemental movie 2**. Video showing representative synthetic images for each input *Y→X* Spatial proximity from main figure 6 panels C and D.

**Figure S1.**
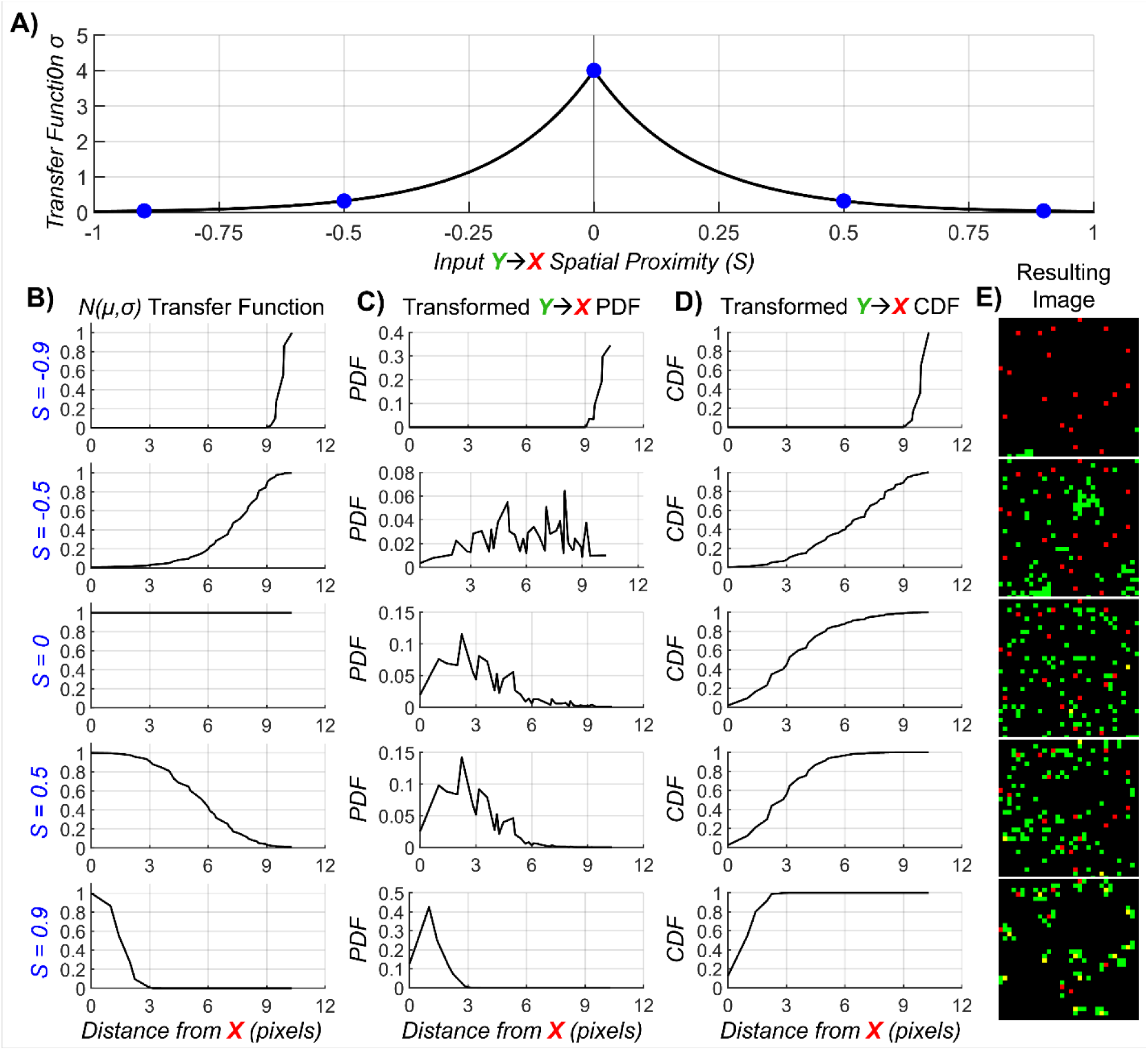
(**A**) Empirical function, defined by x-coordinates linearly spaced between 0 and 1 and y-coordinates logarithmically spaced between 0.026 and 4, to convert the input *Y→X* spatial proximity parameter *S* into a standard deviation σ value for the half normal distribution *N(μ,σ)* used for transforming the *Y→X* distance assignment-PDF. Blue dots indicate sample *S* and *σ* values used for generating (**B**) exemplar *N(μ,σ)* PDF transfer functions. (**C**) *N(μ,σ)*-transformed PDFs defining the distribution of distances between *Y*-events and their closest *X*-events. (**D**) The CDFs derived from the (**C**) PDFs used to assign each *Y*-event a distance-to-*X* using the inverse-CDF method. (**E**) Resulting synthetic images where the mean distance between *Y*-events (green) and *X*-events (red) have been tuned using the parameter *S*. (**B middle**) When *S* is 0, the *N(μ,σ)*-transfer function is equal to 1 at all distance, causing the assignment-PDF to be unaltered and *Y*-events to be assigned distances expected under CSR. (**B top**) When *S* is negative, the *N(μ,σ)*-transfer function is biased to favor larger distances, thus increasing their frequency in the assignment-PDF and causing *Y*-events to generally be assigned distances greater than expected under CSR. (**B bottom**) When *S* is positive, the *N(μ,σ)*-transfer function is biased to favor smaller distances, thus increasing their frequency in the assignment-PDF and causing *Y*-events to generally be assigned distances smaller than expected under CSR.

**Figure S2.**
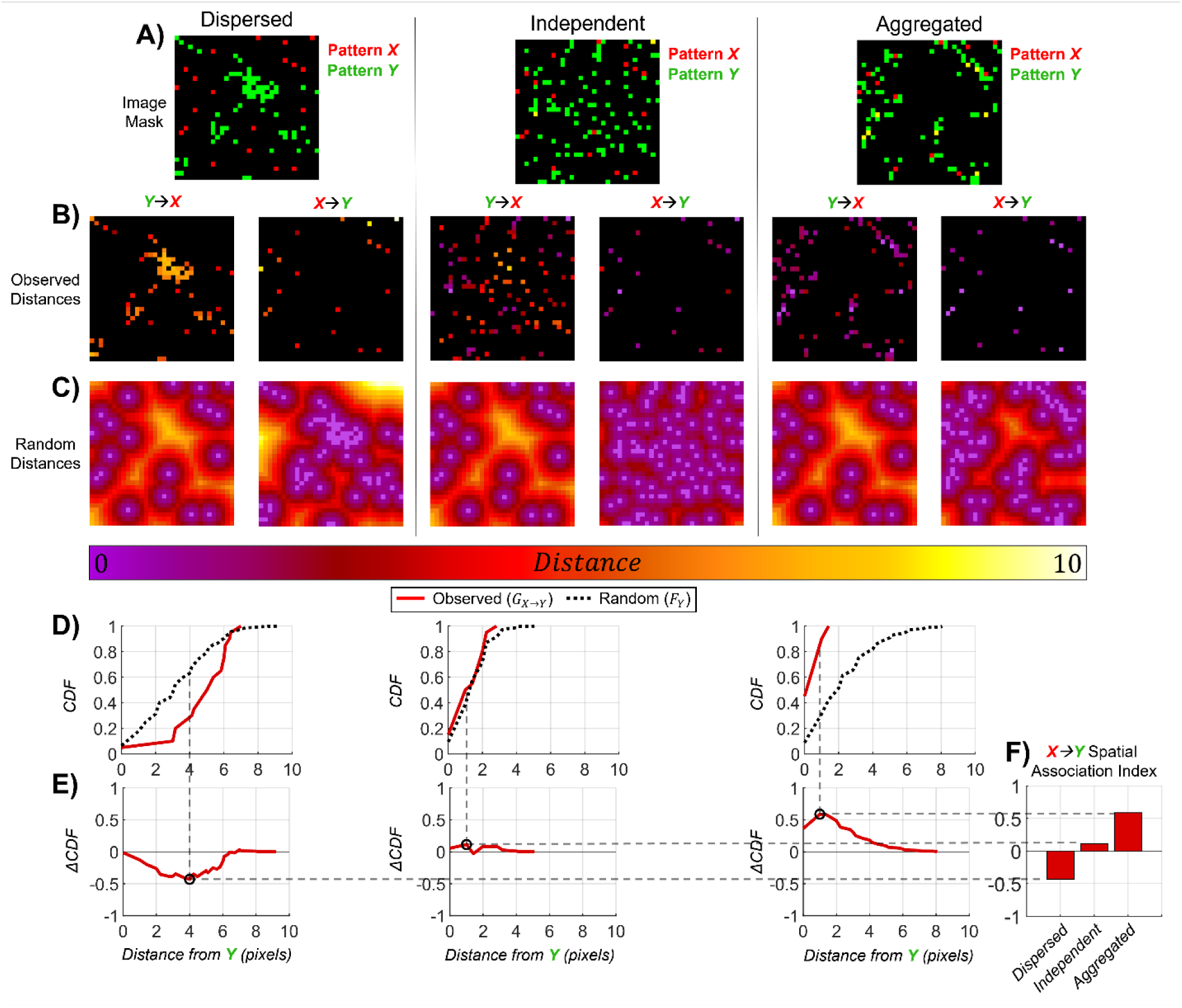
(**A**) 2D synthetic images exhibiting (**A left**) dispersed, (**A middle**) independent, and (**A right**) aggregated patterns. (**B**) Heatmaps of observed distances between each subject-event and the nearest landmark-event. (**C**) Heatmaps of random distances that are the distances between every pixel in the image and the nearest landmark-event. (**D**) Empirical CDFs of *X→Y* distances measured from panel **A** images. Observed CDFs are plotted with solid red lines and random CDFs are plotted with dotted black lines. (**E**) Delta CDFs representing deviation from CSR as a function of distances with black circles marking their peaks. (**E**) Resulting spatial association indices with asterisks indicating significant deviation from CSR evaluated using a 2-sided KS test. Horizontal and vertical dashed gray lines respectively indicate the values and distances of delta function peaks from which spatial association indices were derived.

**Figure S3.**
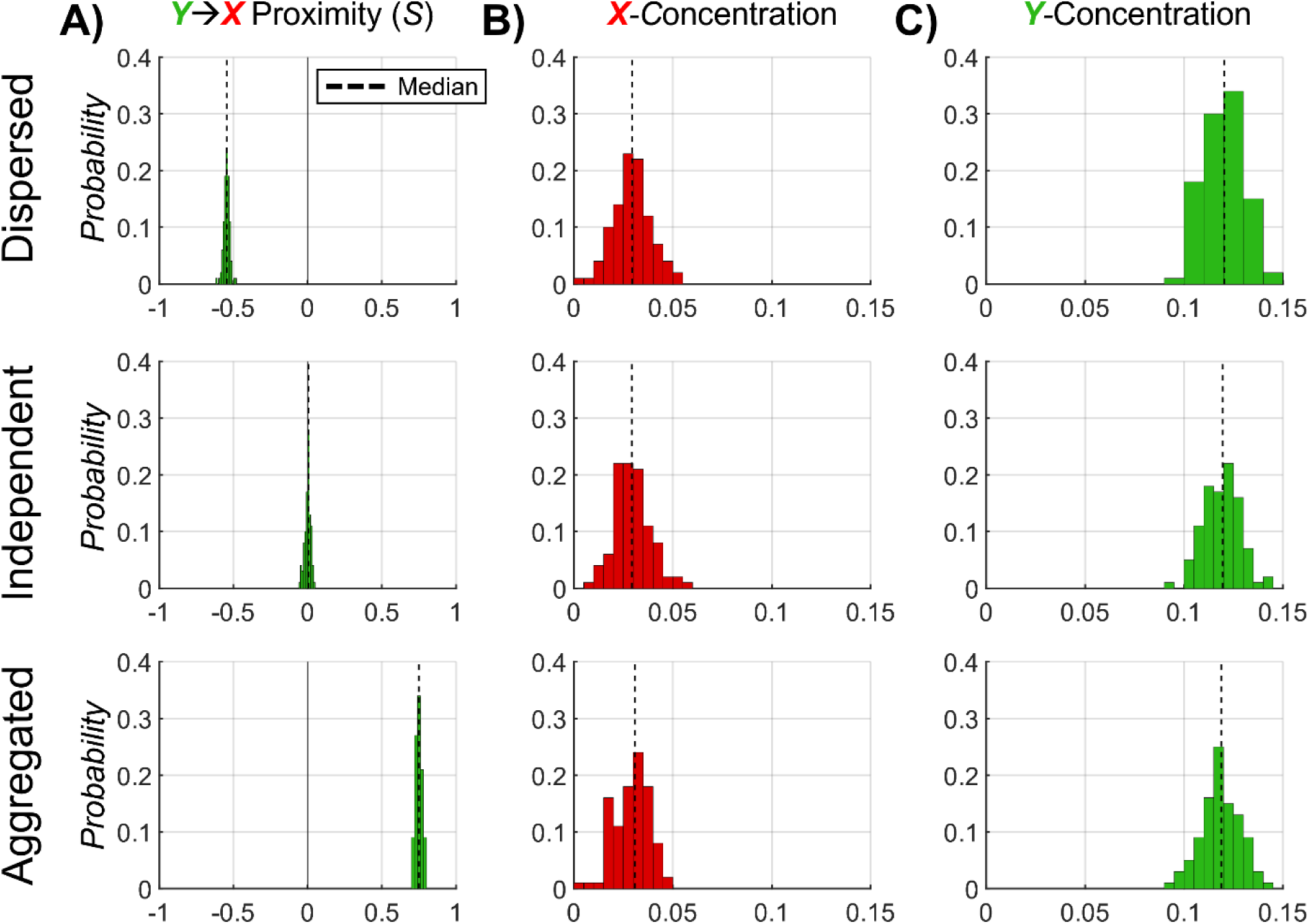
Histograms for each image group of (**A**) input *Y→X* spatial proximity parameter (*S*), (**B**) pattern-X concentration, and (**C**) pattern-Y concentration, used to generate the synthetic from main figure 3 and supplemental figure 4. Dashed lines indicate distribution medians.

**Figure S4.**
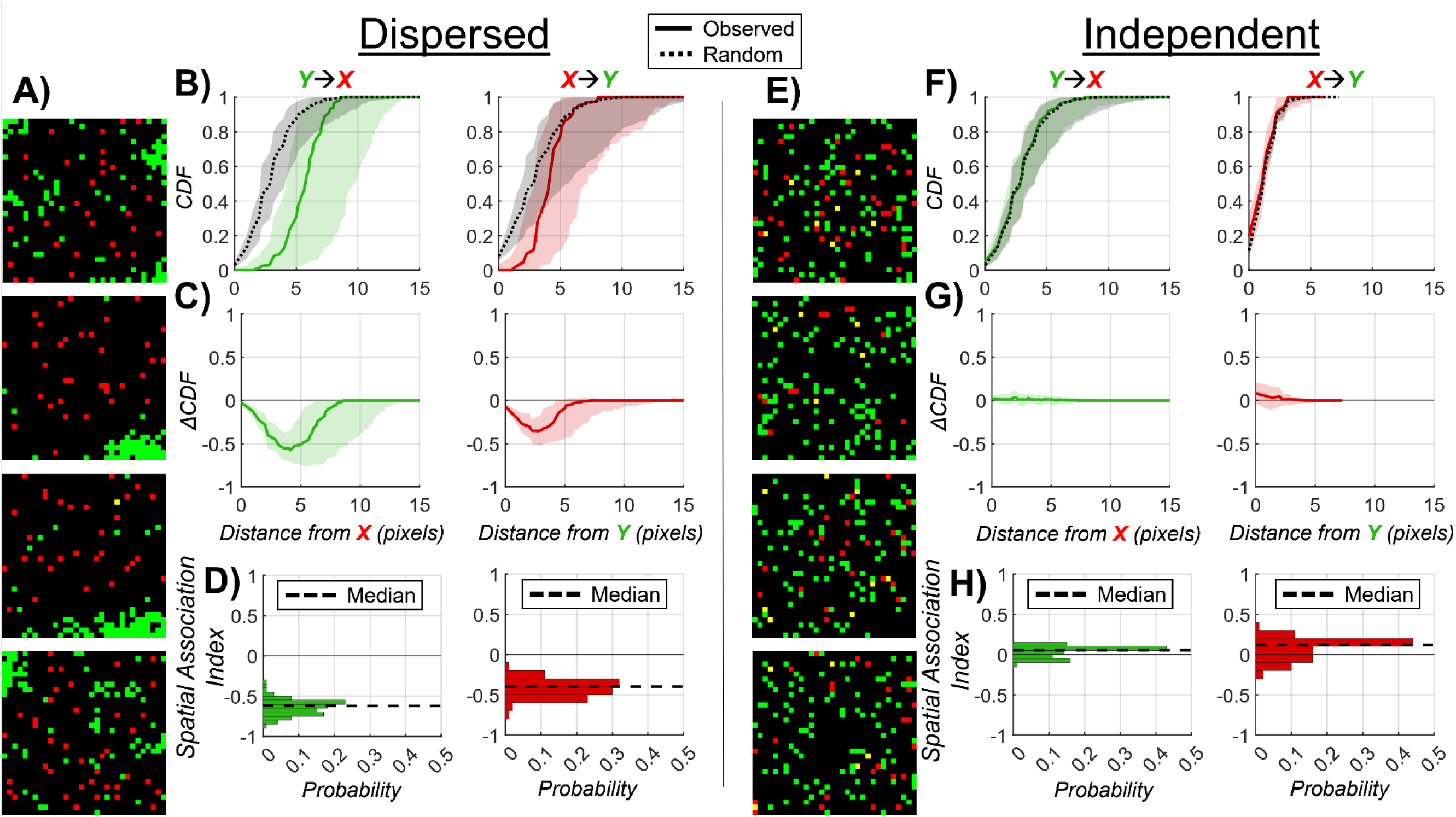
(**A, E**) Representative 2D synthetic images exhibiting (**A**) dispersion and (**E**) independence between pattern-*X* (red) and pattern-*Y* (green) patterns. (**B, F**) Weighted median CDFs and (**C, G**) delta CDFs for both perspectives of SPACE analysis. Shaded regions indicate upper 95% and lower 5% weighted quantiles. (**B, F**) Observed CDFs are plotted with solid lines and random CDFs with dashed lines. (**D, H**) Histogram of *Y→X* (green) and *X→Y* (red) spatial association indices measured from the entire batch of synthetic images with spatially (**D**) dispersed and (**H**) independent patterns. Dashed black lines indicate distribution medians. Each group was composed of 100 images.

**Figure S5.**
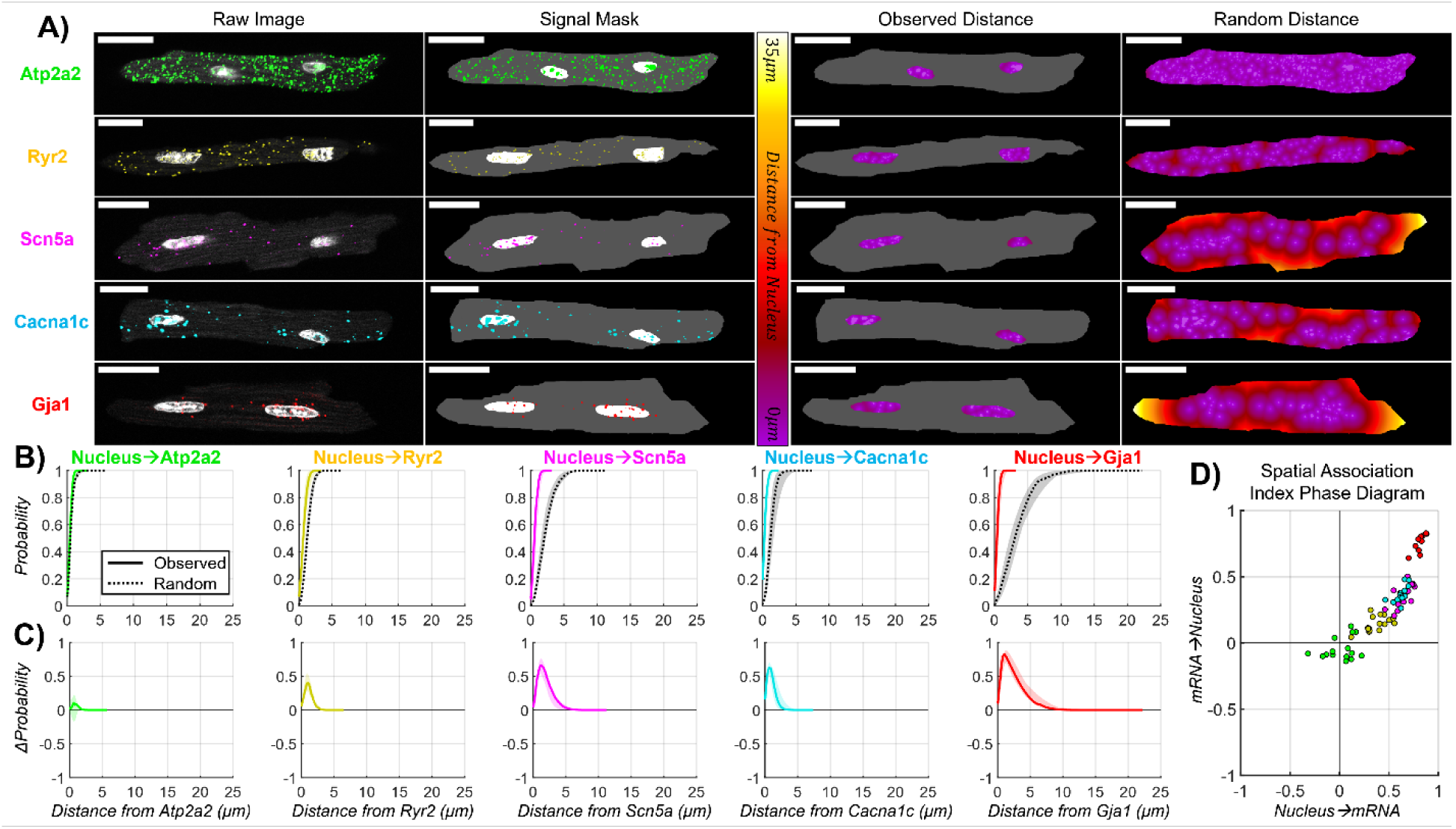
(**A left**) Representative 2D planes from 3D confocal images of individual cardiomyocytes with fluorescently labeled nuclei (white) and mRNAs Atp2a2 (green), Ryr2 (yellow), Scn5a (magenta), Cacna1c (cyan), and Gja1 (red). Scale bars are 20μm. (**A middle-left**) Image masks indicating event pixels for mRNA (colors) and nuclei (white) and the cell body as the study area (grey). (**A middle-right**) heatmaps of observed distances between each nucleus mask pixel and their closest mRNA mask pixel. (**A right**) heatmaps of random distances that are the distances between each pixel in the study area and their closest mRNA pixel. (**B**) Weighted median CDFs and (**C**) delta CDFs for the *Nucleus→mRNA* SPACE analysis. Shaded regions indicate upper 95% and lower 5% weighted quantiles. (**B**) Observed CDFs are plotted with solid lines and random CDFs with dashed lines. (**D**) Spatial association index phase diagram plotted as the bivariate histogram of *Nucleus→mRNA* spatial association indices for each image on the x-axis and *mRNA→Nucleus* spatial association indices on the y-axis. *mRNA→Nucleus* spatial association indices were compared between mRNAs using Bonferroni-corrected pairwise weighted Student’s T-tests (**Supplemental Table 2**). (Gja1: n = 10 cells from 3 hearts; Scn5a: n = 16 cells from 3 hearts; Cacna1c: n = 16 cells from 3 hearts; Ryr2: n = 16 cells from 3 hearts; Atp2a2: n = 15 cells from 3 hearts).

**Figure S6.**
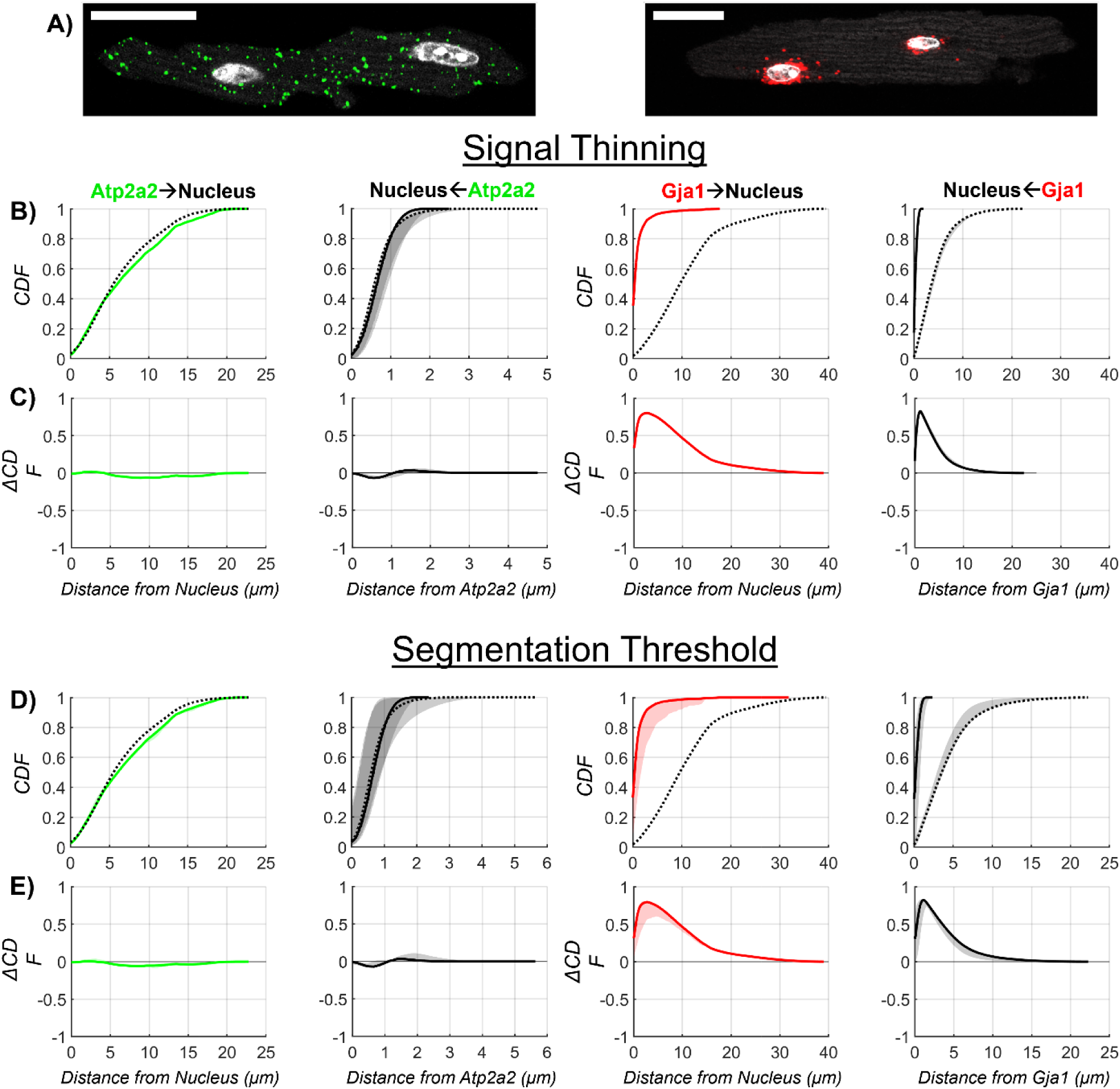
(**A**) 2D planes from 3D confocal images of individual cardiomyocytes with fluorescently labeled nuclei (white), Atp2a2 mRNA (green), and Gja1 mRNA (red). Scale bars are 20μm. (**B, D**) Weighted median CDFs and (**C, E**) delta CDFs showing the central tendency and spread of functions for all (**B, C**) thinning and (**D, E**) segmentation intensity threshold values evaluated in main figures 5 and 6. Shaded regions indicate maximum and minimum function values at each distance. (**B**) Observed CDFs are plotted with solid lines and random CDFs with dashed lines.

**Table S1.** p-values for the Bonferroni-correct pairwise weighted Student’s T-tests comparing the (**A**) *X→Y* and (**B**) *Y→X* spatial association indices between image groups for synthetic data from main figure 2 and supplemental figure 4.

| **A)** $X \rightarrow Y$ Spatial Association Index p-values | | | |
| --- | --- | --- | --- |
|  | Dispersed | Independent | Aggregated |
| Dispersed | 2 | 0.0060 | 4.08E-10 |
| Independent | 5.979E-03 | 2 | 0.0141 |
| Aggregated | 4.077E-10 | 0.0141 | 2 |

**Table S1.** p-values for the Bonferroni-correct pairwise weighted Student's T-tests comparing the (A) $X \rightarrow Y$ and (B) $Y \rightarrow X$ spatial association indices between image groups for synthetic data from main figure 2 and supplemental figure 4.
| **B)** $Y \rightarrow X$ Spatial Association Index p-values | | | |
| --- | --- | --- | --- |
|  | Dispersed | Independent | Aggregated |
| Dispersed | 2 | 1.762E-07 | 8.86E-16 |
| Independent | 1.762E-07 | 2 | 3.252E-05 |
| Aggregated | 8.858E-16 | 3.252E-05 | 2 |

**Table S2.** p-values for the Bonferroni-correct pairwise weighted Student’s T-tests comparing the (**A**) *mRNA→Nucleus* and (**B**) *Nucleus→mRNA* spatial association indices between the different mRNAs from main figure 3 and supplemental figure 5.

| mRNA→Nucleus Spatial Association Index p-values |  |  |  |  |  |
| --- | --- | --- | --- | --- | --- |
| **A)** | **Atp2a2** | **Cacna1c** | **Gja1** | **Ryr2** | **Scn5a** |
| **Atp2a2** | 4 | 0.0016 | 3.2689E-06 | 0.3633 | 0.0136 |
| **Cacna1c** | 0.0016 | 4 | 0.0021 | 0.0317 | 3.2103 |
| **Gja1** | 3.2689E-06 | 0.0021 | 4 | 1.8716E-06 | 0.0120 |
| **Ryr2** | 0.3633 | 0.0317 | 1.8716E-06 | 4 | 0.2037 |
| **Scn5a** | 0.0136 | 3.2103 | 0.0120 | 0.2037 | 4 |

| Nucleus→mRNA Spatial Association Index p-values |  |  |  |  |  |
| --- | --- | --- | --- | --- | --- |
| **B)** | **Atp2a2** | **Cacna1c** | **Gja1** | **Ryr2** | **Scn5a** |
| **Atp2a2** | 4 | 0.0059 | 0.0038 | 0.4152 | 0.0078 |
| **Cacna1c** | 0.0059 | 4 | 0.2018 | 0.2612 | 3.1149 |
| **Gja1** | 0.0038 | 0.2018 | 4 | 0.0419 | 0.9399 |
| **Ryr2** | 0.4152 | 0.2612 | 0.0419 | 4 | 0.2626 |
| **Scn5a** | 0.0078 | 3.1149 | 0.9399 | 0.2626 | 4 |

