## Supplementary figures and images for "Spatial Pattern Analysis using Closest Events (SPACE) – A Nearest Neighbor Point Pattern Analysis Framework for Assessing Spatial Relationships from Image Data"

### Supplemental Figure 1

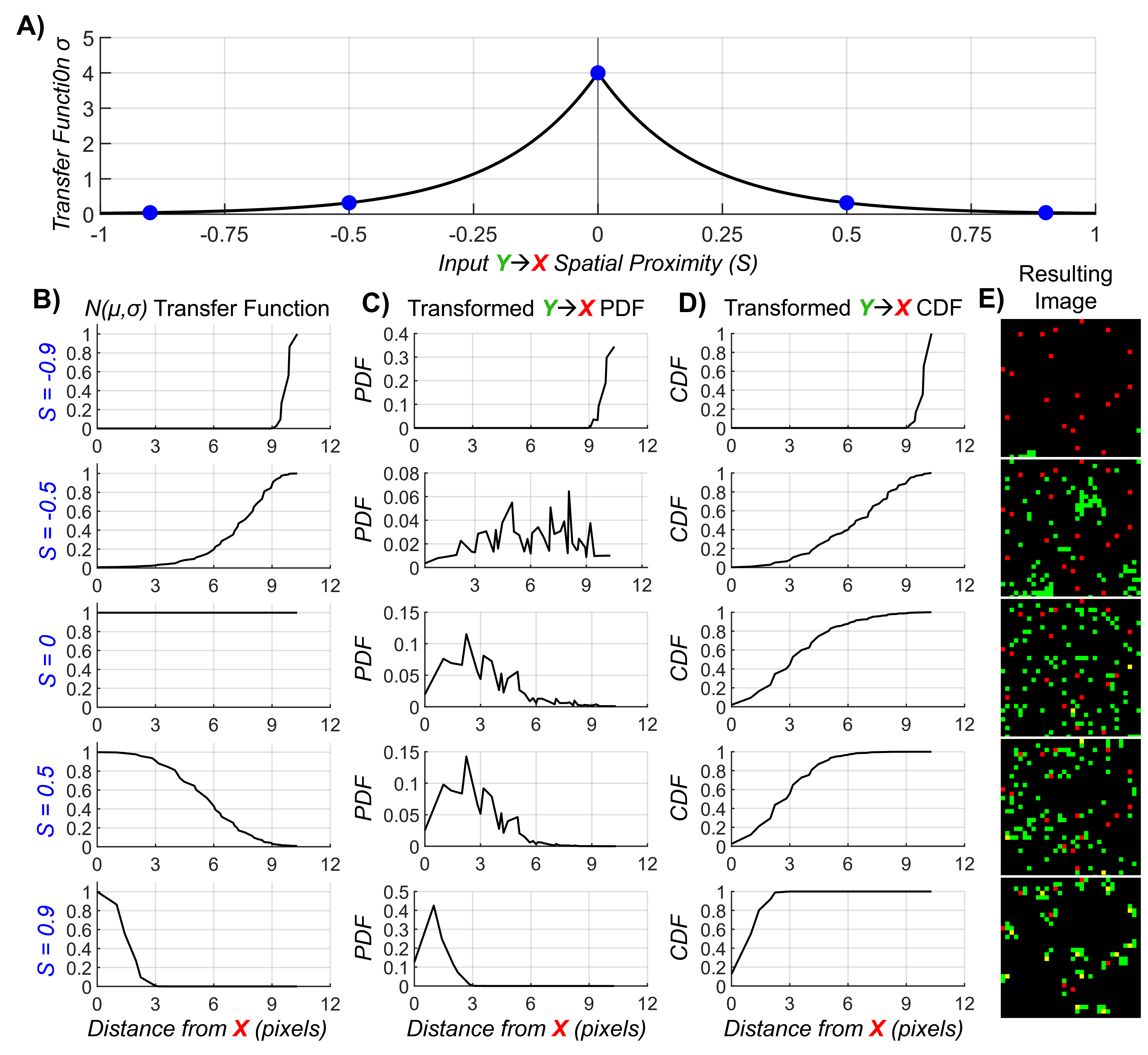

### Supplemental Figure 2

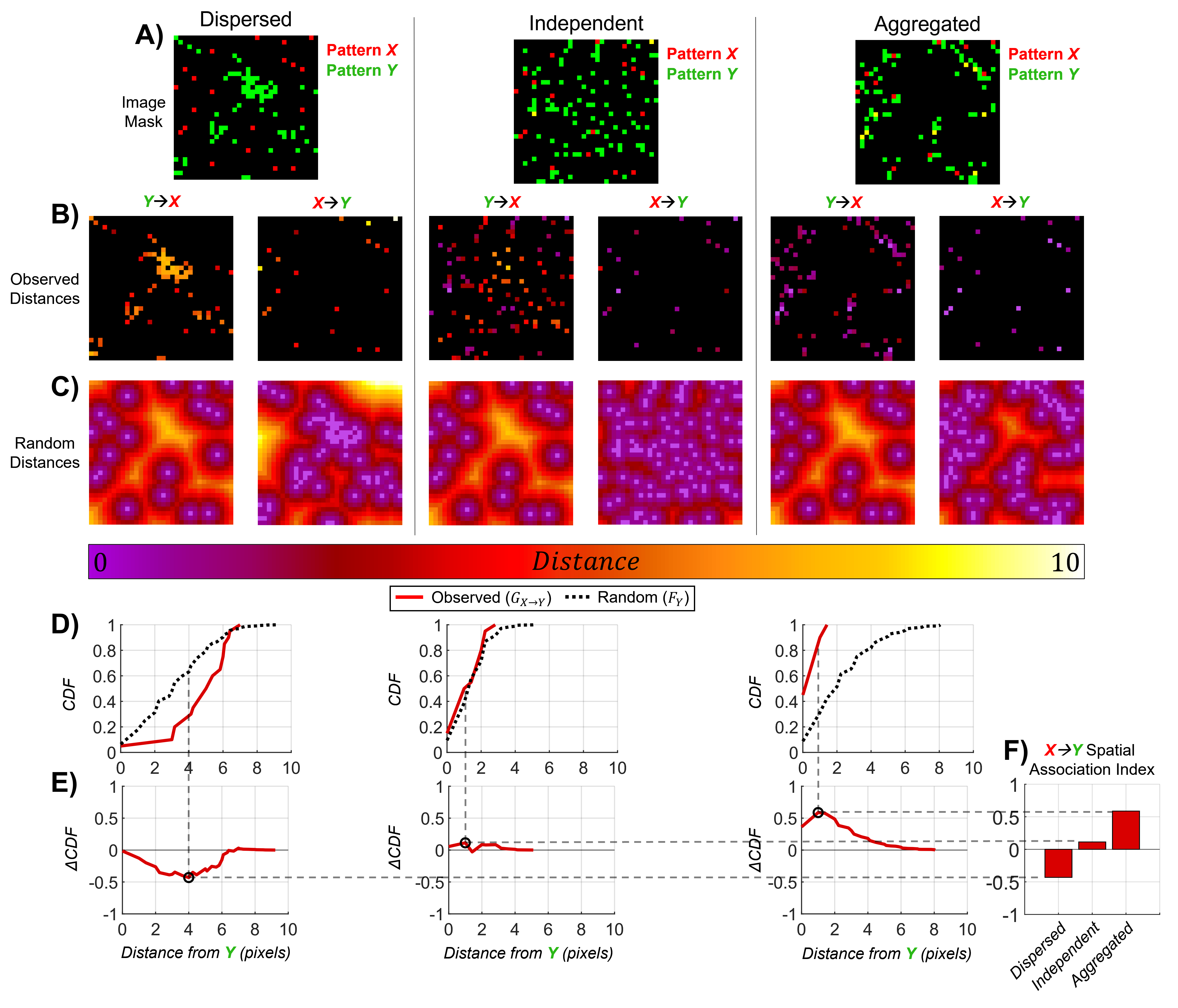

### Supplemental Figure 3

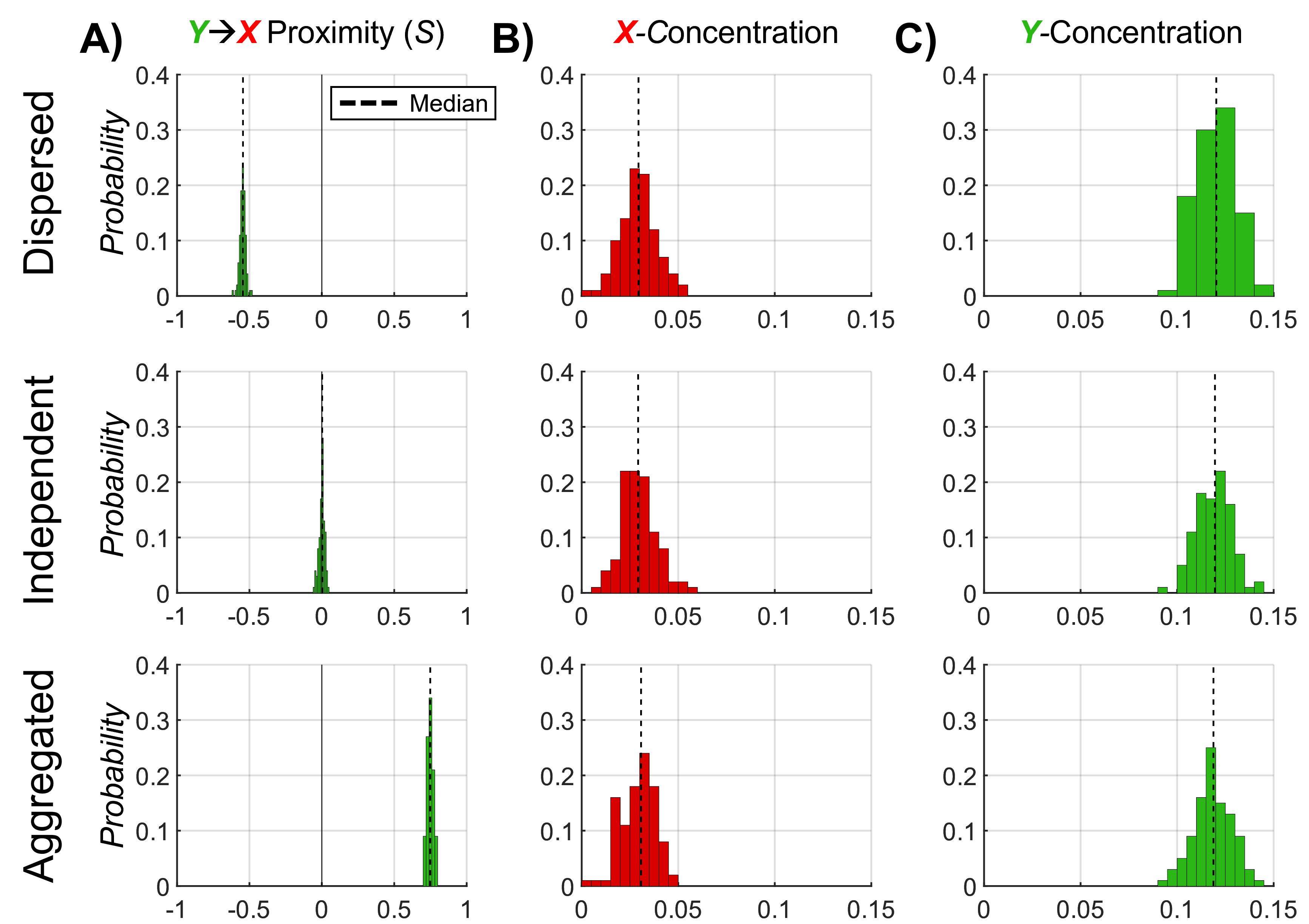

### Supplemental Figure 4

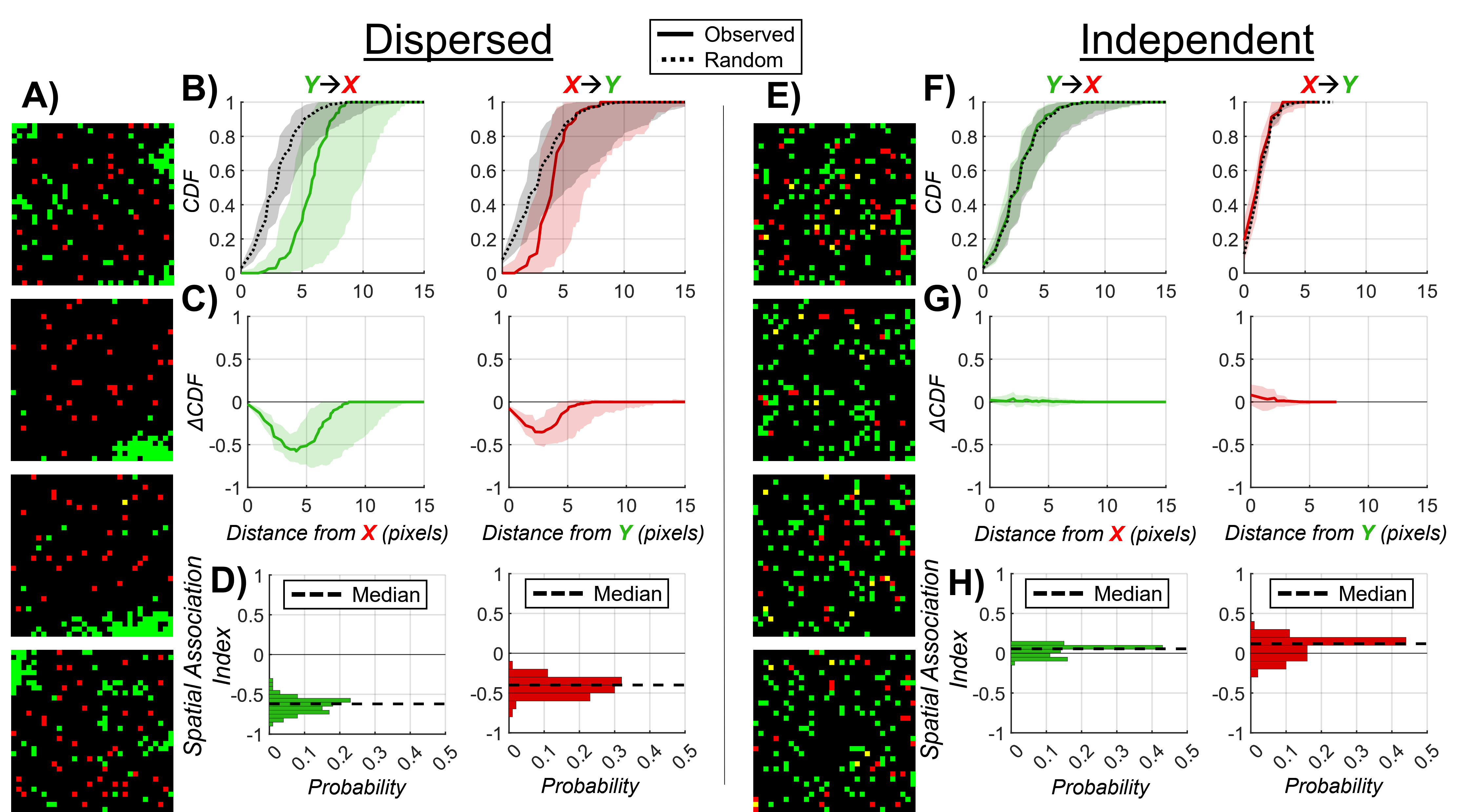

### Supplemental Figure 5

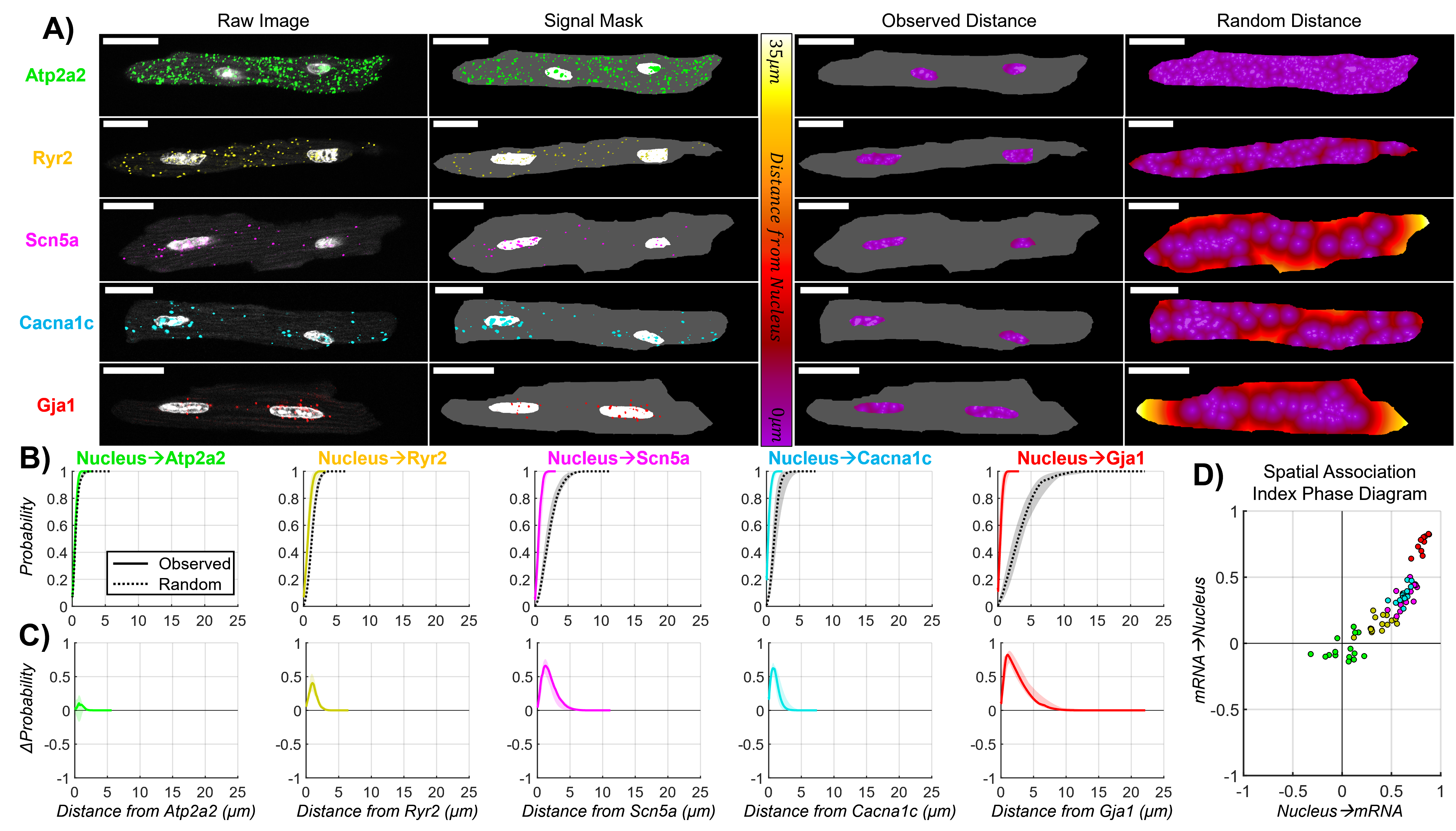

### Supplemental Figure 6

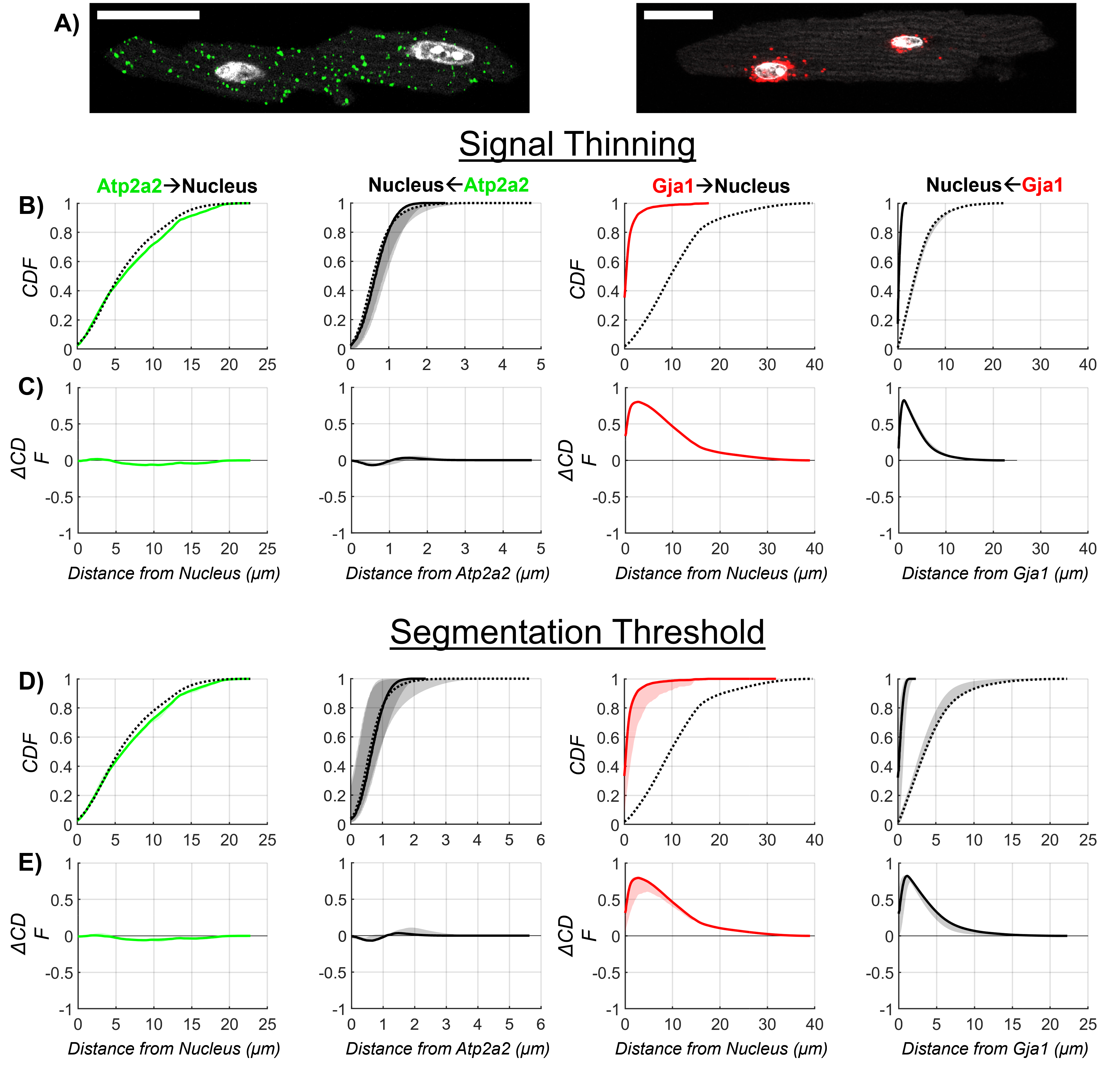

### Supplemental Table 1

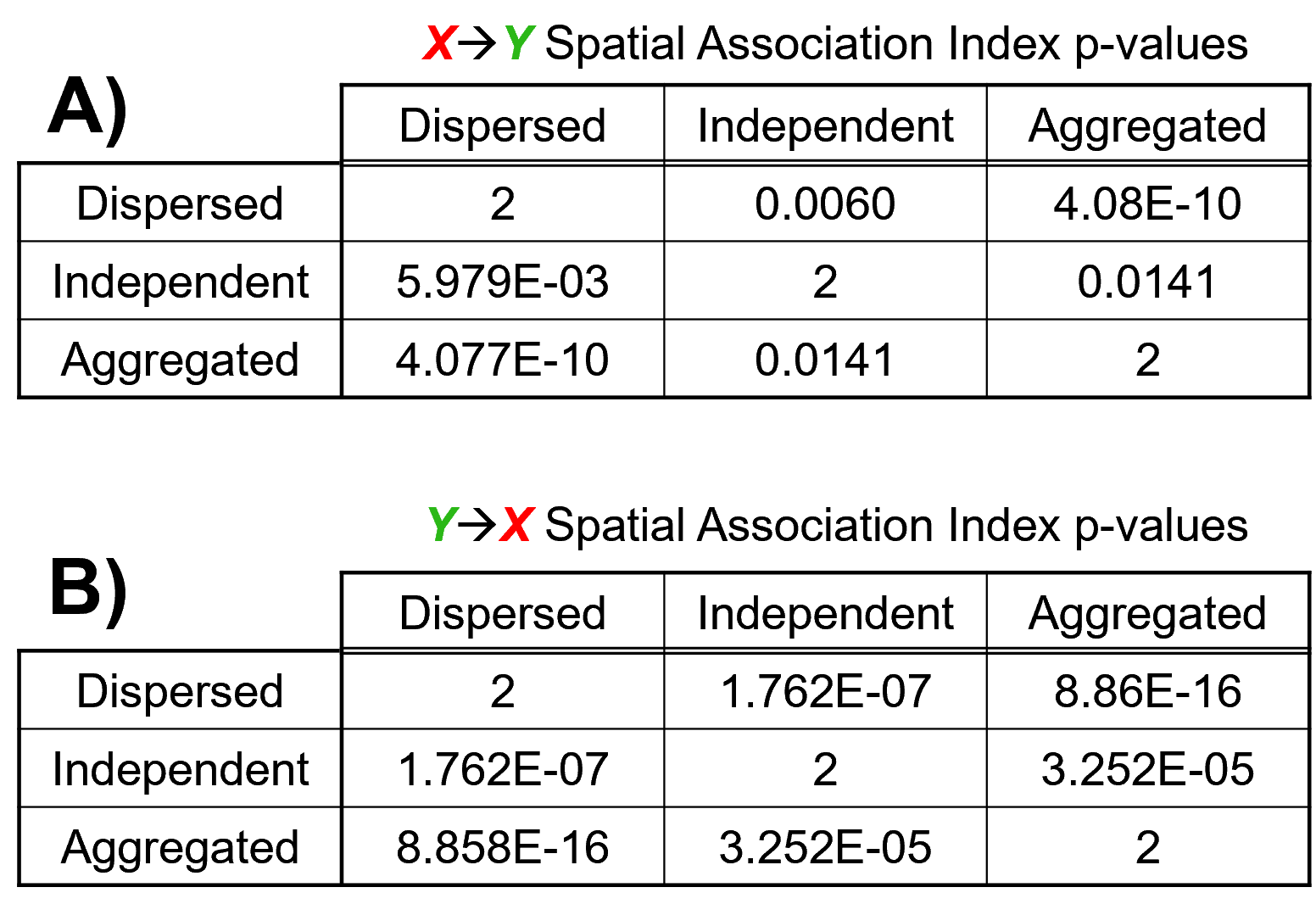

### Supplemental Table 2

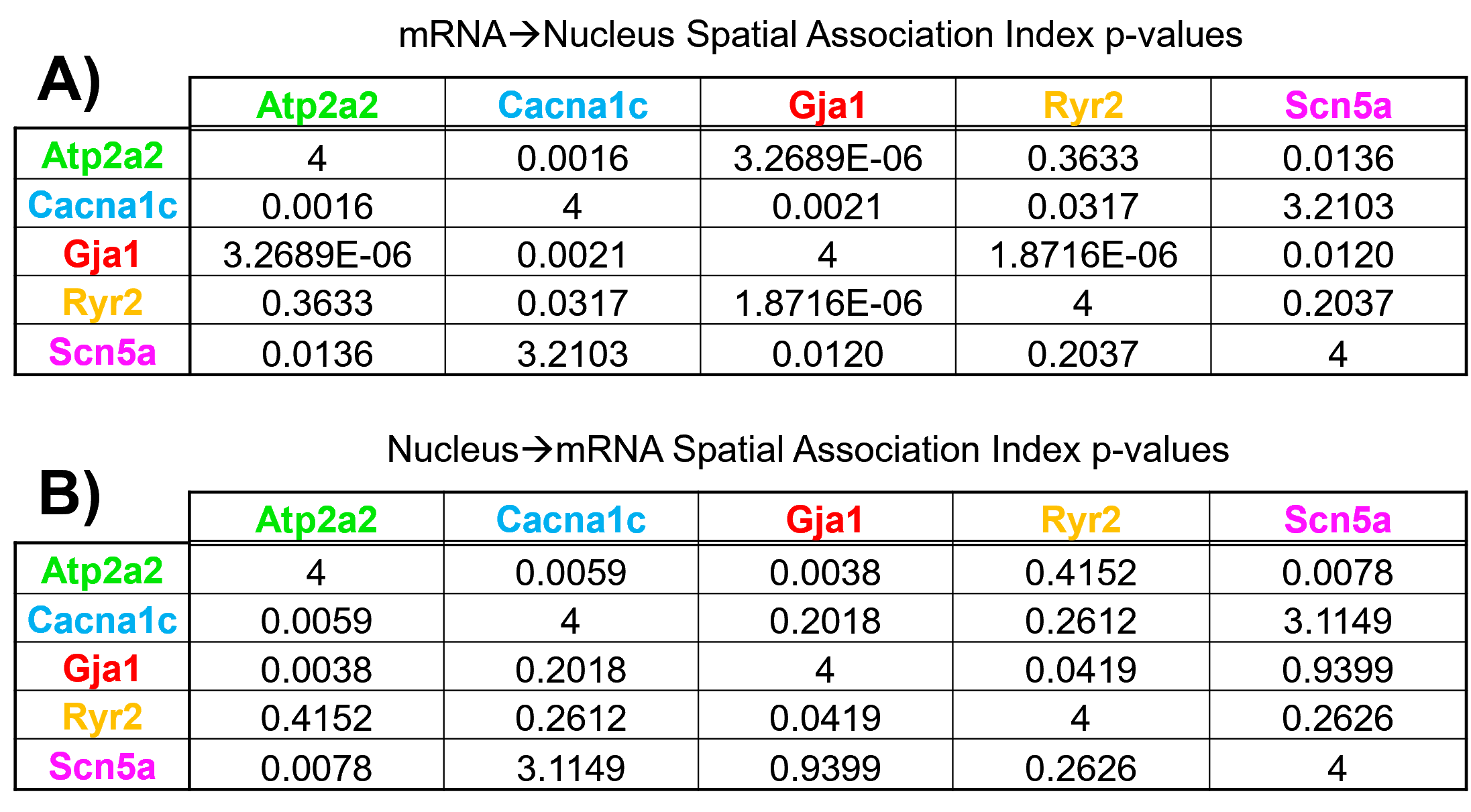
